# A Human Next Generation PBK Model for PFOA

**DOI:** 10.64898/2026.02.04.703497

**Authors:** Chrysanthi Pachoulide, Carolina Vogs, Aude Ratier, Jack Koster, Trine Husøy, Martine Vrijheid, Yiyi Xu, Antonios Georgelis, Karl Forsell, Joost Westerhout, Nynke I. Kramer

**Author notes:** Corresponding author’s.

## Abstract

The human toxicological risk assessment of per- and polyfluoroalkyl substances (PFAS) is challenging, due to their sheer number and structural diversity, but also the paucity of the toxicity data required to characterize them. The development of Next Generation Physiologically Based Kinetic (NG-PBK) models may assist in overcoming this challenge. The mechanistic nature of NG-PBK models allows for their extrapolation from data-rich PFAS, such as perfluorooctanoic acid (PFOA), to data-poor ones, facilitating their application in Next Generation Risk Assessment (NGRA). The present study proposes a NG-PBK model for PFOA in humans, parametrized exclusively using in vitro-, and in silico-derived data. The model describes the toxicokinetic processes of 1) partitioning to plasma and tissue proteins, 2) partitioning to cell membrane lipids, 3a) transporter-mediated entero-hepatic circulation and 3b) renal elimination and reabsorption, and 4) elimination via menstruation. Global sensitivity analysis indicated that the model was most sensitive to the fraction unbound in plasma, active-transport parameters, and tissue-plasma partition coefficients. The model was equivalent to already available validated human PFOA-PBK models, while compared to those, it is not calibrated to observed animal, nor human data, illustrating its strength in being mechanistic. The serum concentrations and half-lives predicted by the NG-PBK model were within the ranges of those reported in human volunteer and biomonitoring (HBM) studies, demonstrating the model’s capacity to accurately predict PFOA toxicokinetics on exposure estimates. Extrapolation of the NG-PBK model to other PFAS, in conjunction with its integration with HBM data, will facilitate the NGRA of PFAS. This is particularly relevant given the paucity of in vivo data for most PFAS, ensuring compliance with the 3R principles.

## 2 Introduction

Human exposure to per- and polyfluoroalkyl substances (PFAS), a class of thousands of prevalent environmental pollutants of anthropogenic nature, poses a risk to human health. The increasing evidence that PFAS exposure is associated with adverse health effects has led to the regulation of legacy PFAS, including perfluorooctanoic acid (PFOA), perfluorooctane sulfonic acid (PFOS), perfluorononanoic acid (PFNA) and perfluorohexane sulfonic acid PFHxS, but also the increase in production and emission of alternative and precursor PFAS. To this date, legacy PFAS and more than 40 PFAS alternatives and precursors have been detected in human blood matrices in human biomonitoring (HBM) studies (Borghese et al., 2024; Schultz et al., 2023). Consequently, the whole PFAS class poses a potential risk to human health and hence, is a priority in human toxicological risk assessment (Rosato et al., 2022; Fenton et al., 2020). Humans are exposed to PFAS mainly through ingestion, although dermal absorption and inhalation also occur (Richterová et al., 2023; Bil et al., 2022; Sunderland et al., 2018; Poothong et al., 2020).

Exposure to PFAS has been associated with adverse health effects, including immunosuppression (Crawford et al., 2023; Hong et al., 2025; Huang et al., 2024; Abraham et al., 2020), cardiovascular, metabolic, endocrine, reproductive and developmental toxicity, diseases of the nervous system (Radke et al., 2022; Shirke et al., 2024), and cancer (Biggeri et al., 2024; Li et al., 2025). The risk of developing adverse health effects not only depends on the hazardous properties of PFAS, but also on their toxicokinetics in the human body, *i*.*e*., absorption, distribution, metabolism and elimination. Toxicokinetic processes drive the relationship between external and internal exposure in different body parts, including the systemic blood circulation and target tissue(s) of toxicity. Consequently, integration of HBM, toxicokinetics and toxicity data increases the weight of evidence used for their toxicological risk assessment. PFAS risk assessment following the traditional one-by-one, laboratory animal-based, toxicity testing approach is challenging due to the sheer number (> 4700 (OECD, 2021b)) of PFAS.

A potential solution to this challenge lies in the implementation of the Next Generation Risk Assessment (NGRA) framework. NGRA does not rely on laboratory animal tests but promotes the application of new approach methodologies (NAMs), in a tiered approach. NAMs aim to provide a mechanistic understanding of the toxicity and toxicokinetics, while being high-throughput and alternatives to animal testing (Dent et al., 2018; Paini et al., 2021a,b). For instance, mechanistic NAM studies demonstrated that the toxicokinetics of PFAS are driven by their binding to plasma proteins and cell-membrane phospholipids (Fischer et al., 2025; Allendorf et al., 2020; Zhao et al., 2023). The capacity of PFAS to occupy the binding cavity of proteins, as well as their affinity to cell-membrane phospholipids, are both influenced by the number of per-fluorinated carbons (npc), and their functional group. The understanding of these toxicokinetic properties, and structure-property relationships, obtained via NAMs, can facilitate the hazard characterization of PFAS, thus circumventing the necessity for one-by-one laboratory animal tests.

One of these NAMs is physiologically-based kinetic (PBK) modelling. PBK models are mathematical models that simulate the fate of a chemical in an organism, based on the physicochemical and biochemical properties of the chemical and the physiology of the organism studied. Several human-PBK models are available for a limited number of PFAS, mainly PFOA and PFOS, but they were generally calibrated to in vivo toxicokinetic data. (Loccisano et al., 2011; EFSA, 2020; Cheng and Ng, 2017; Lin et al., 2023; Ratier et al., 2024; Husøy et al., 2023; Karakoltzidis et al., 2025). These models need to be calibrated to in vivo data in order to correct for discrepancies between predicted and observed toxicokinetic properties/profiles (maximum concentration (Cmax), area under the concentration-time curve (AUC), half-life). These discrepancies may be attributed to a poor, or incomplete depiction of key biokinetic processes, such as transporter mediated uptake and elimination in the liver and kidney and binding to tissue constituents, both of which influence the biodistribution of PFAS. A consequence of calibrating/fitting, is that extrapolation of these traditional PBK models to other PFAS congeners requires the calibration of each model on congener-specific in vivo data.

A way forward might be the development of next generation PBK (NG-PBK) models. The structure of NG-PBK models is informed by the knowledge of the chemical-specific biokinetic mechanisms (such as protein-binding, metabolism, active transport) obtained from NAM data (OECD, 2021a). By being mechanistic, these models provide a physiologically and biologically plausible explanation of the toxicokinetic behaviour of a chemical, therefore overcoming the need of calibration to animal or human data (Paini et al., 2019). For instance, by refining the structure of a generic PBK model to include transporter-mediated cellular uptake and elimination of benzophenone-4 in the liver and kidney, (Punt et al., 2025) was able to predict relevant internal concentrations of the chemical used in its NGRA, without the use of any animal data.

In this study, we aim to develop a mechanistic NG-PBK model that builds on recent advancements in our understanding of membrane, lipid, and protein binding kinetics of PFAS, combined with human physiology. For the development of the NG-PBK model, we focused on PFOA as a proof of concept, but the model can also be applied to other PFAS with similar structural properties in the future, including those lacking in vivo kinetic information (Thompson et al., 2024). To this end, we first developed the modelling framework and upscaled essential toxicokinetic parameter values obtained using NAMs to humans. We then evaluated the developed PBK model against HBM data spanning daily intakes, age-groups, and exposure routes by comparing predicted PFOA plasma concentrations after oral and dermal exposure over a whole lifetime to observed plasma concentrations (Ratier et al., 2024; Johanson et al., 2023; Olsen et al., 2007).

To facilitate the regulatory acceptance of the developed NG-PBK model as suggested by experts and regulatory agencies (Paini et al., 2021b,a), we followed the OECD guidance document, which incorporates the scientific workflow for characterizing, validating and evaluating, including the model’s goodness-of-fit and sensitivity analyses (OECD, 2021a). Lastly, we demonstrated how the incorporation of physiological data can explain the high inter-individual variability in PFOA serum concentration, and half-life in humans observed in HBM studies. We made our PBK tool open accessible for use to facilitate its extrapolation to other PFAS:(Pachoulide, 2025).

## 3 Theoretical toxicokinetic basis for the NG-PBK model development

This study proposes a NG-PBK model that mechanistically incorporated the toxicokinetic processes specific to PFAS based on in vitro and in silico data, including: 1) partitioning to plasma and tissue proteins (Fischer et al., 2024; Ryu et al., 2024; Allendorf et al., 2019, 2020), 2) partitioning to cell membrane lipids (Ebert et al., 2020; Allendorf et al., 2020), and 3) the transporter-mediated processes of entero-hepatic circulation (EHC) and renal elimination and reabsorption (RER) (Kimura et al., 2017; Janssen et al., 2024; Lin et al., 2023). In addition, we added elimination via menstruation (Wu et al., 2015; Upson et al., 2022) as an excretion pathway of PFAS in women.

### 3.1 Partitioning to plasma and tissue proteins

PFOA is assumed to be distributed within the organism based on two factors: plasma perfusion rate of each tissue, and sorption of PFOA to tissue constituents. In blood, PFOA does not partition to red blood cells, but mostly binds to plasma proteins (Ehresman et al., 2007; Fischer et al., 2024). In plasma, PFOA has a high affinity to albumin and to a lesser extent to globulins (Fischer et al., 2024; Ryu et al., 2024; Smeltz et al., 2023). Since PFOA is bound to plasma proteins, plasma loss due to menstrual bleeding contributes to the excretion of PFOA (Wu et al., 2015; Upson et al., 2022). The amount of PFOA available in the systemic plasma circulation, is free to be distributed to different tissues, based on its tissue to plasma partition coefficients 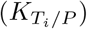.

### 3.2 Partitioning to tissue constituents

Distribution of PFOA to tissue constituents is driven by its sorption affinity to each respective constituent and its fractional volume (Utsey et al., 2020; Allendorf et al., 2020), while excretion of PFOA from the tissue depends on the freely available fraction of PFOA in water (fraction unbound (fu)) (Figure S3). Thus, tissue-to-plasma partition coefficients are estimated by describing each tissue as a combination of the fractional volumes of neutral (storage) lipids, membrane lipids, albumin, structural proteins, fatty acid binding proteins (FABP) and water. Previous studies experimentally determined the sorption affinities of PFOA to these constituents (Ebert et al., 2020; Allendorf et al., 2019, 2020), showing that PFOA exhibited the highest affinity for albumin, FABP and membrane lipids, followed by a lower affinity for structural proteins, and a low affinity for storage lipids (Figure S4). This finding suggested that PFOA tends to accumulate in organs such as the skin, kidneys, adipose and liver which exhibit higher albumin and/or membrane lipid content compared to other tissues (see Table S1) (Allendorf et al., 2020; Utsey et al., 2020).

### 3.3 Transporter-mediated processes

Lungs, liver, and kidney human autopsy samples had indeed the highest PFOA concentrations compared with other organs (Nielsen et al., 2023). Nonetheless, the 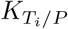 and high perfusion of these tissues alone could not explain the high measured PFOA concentrations and contradict the low concentrations in the brain, suggesting that active transport could enhance the distribution of PFOA in perfusion-rich organs and limit its distribution to the brain (Nielsen et al., 2023). Multiple in vitro and in silico studies investigated the affinities of PFOA to different active transporters (Figure S8) that potentially contribute to the absorption and/or elimination of PFOA in humans (Louisse et al., 2022, 2024; Ryu et al., 2024; Vujic et al., 2024; Cao et al., 2022; Niu et al., 2023; Tiburtini et al., 2024; Zhao et al., 2023; Lin et al., 2023; Kimura et al., 2017; Yang et al., 2010). Figure S8 illustrates all known apical and basolateral transporters found to potentially contribute to the uptake in, or efflux out from enterocytes, hepatocytes, and proximal tubule cells.

#### 3.3.1 Transporter-mediated entero-hepatic circulation

Molecular docking studies suggested that PFOA mimics bile acids (such as taurocholate) and could undergo enterohepatic circulation via the same transporters (i.e., sodium/bile acid co-transporter (*NTCP*), apical sodium dependent bile acid transporter (*ASBT*), bile acid export pump (*BSEP*)) (Cao et al., 2022). However, to date, only the PFOA transport by *NTCP* was studied in vitro, by (Lin et al., 2023), who reported a low affinity of PFOA towards the transporter. Lin et al. further demonstrated that both organic anion transporting proteins (*OATPs) 1B1 (OATP1B1)* and *1B3 (OATP1B3)* increased the hepatic uptake of PFOA in vitro. It is therefore more likely that basolateral *OATP1B1* and *OATP1B3*, and not *NTCP*, contribute to the uptake of PFOA to the hepatocytes. Similarly, PFOA uptake in enterocytes was shown to be mediated by *OATP* transporters, and more specifically *OATP2B1* (Kimura et al., 2017; Lin et al., 2023). To the best of our knowledge, there are no in vitro studies that investigated the excretion of PFOA from hepatocytes to bile. Nonetheless, biliary excretion is primarily driven by active transport in humans, hence it is assumed that this is mediated by BSEP (Dawson et al., 2006).

#### 3.3.2 Transporter-mediated renal elimination and reabsorption

Renal excretion of PFOA is influenced by glomerular filtration (GF) and active transport via apical and basolateral organic anion transporters *(OATs)* responsible for reabsorption (RER) to the systemic circulation. GF is a physical process influenced by the difference in the hydraulic pressure between the glomerular capillaries and the glomerular space (ICRP, 2002). The PFOA amount filtered from the glomerular capillaries is limited by the fu in plasma. Upon disposition of PFOA to the primary urine, or the efferent arteries (plasma leaving the glomerulus), a fraction of PFOA can be taken up to the proximal tubule cells by *OAT4* and *URAT1*, and *OAT1* and *OAT3, respectively*, as demonstrated in in vitro studies (Louisse et al., 2024; Lin et al., 2023; Louisse et al., 2022). From all these transporters, *OAT4* could be the major renal transporter, due to both the highest affinity of PFOA towards the transporter, and the higher fu of PFOA in primary urine compared to plasma given the differences in albumin concentrations (Tojo and Kinugasa, 2012). Consequently, PFOA reabsorption via OAT4 could be contributing to the long half-life of PFOA in humans (Louisse et al., 2024).

Overall, it can be concluded that both the EHC and RER of PFOA are active transport mediated. Figure 1 illustrates the mechanisms that are described in the NG-PBK model presented in this study. To best account for these mechanisms in the NG-PBK model, the final model included the plasma, intestinal, liver, kidney, and adipose tissues with or without a skin compartment depending on the exposure route. Arterial and venous blood were lumped into a single homogeneous blood compartment, because heart was not an explicitly modelled organ (Li and Zhang, 2023). Additionally, we assumed that plasma and not blood was the main circulatory system, because PFOA is only distributed in the plasma fraction of blood (Ehresman et al., 2007; Fischer et al., 2024). Lastly, it was assumed that plasma and serum concentrations were equal, because serum to plasma ratios of PFOA were found to be 1:1 (Ehresman et al., 2007).

**Figure 1.**
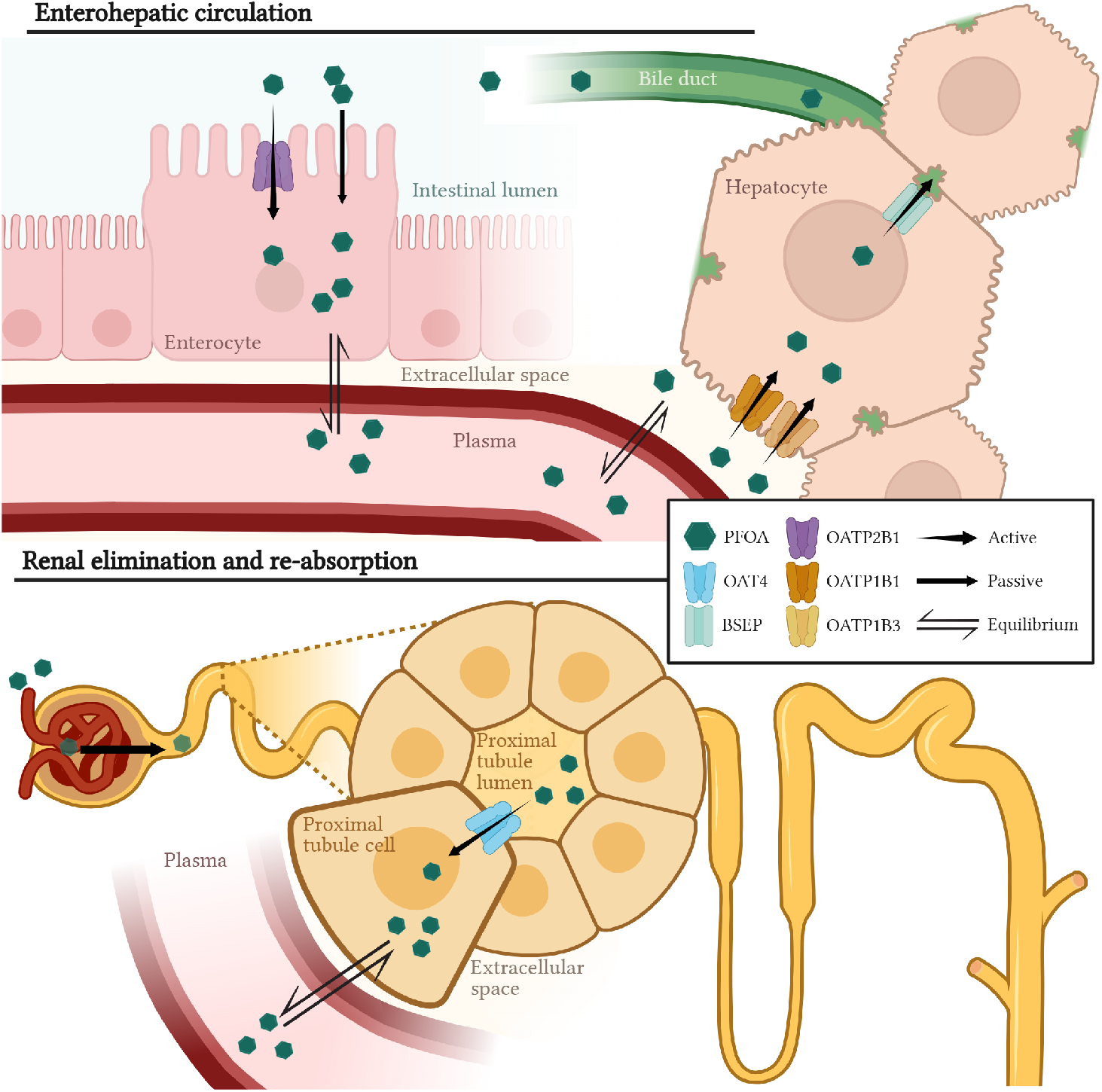
A graphical illustration of A) intestinal absorption and enterohepatic circulation processes, and B) renal excretion and re-absorption of PFOA, as mechanistically applied in the NG-PBK model. Arrows indicate the directions of PFOA mass transfer. A) PFOA is absorbed in the enterocytes by both passive diffusion and active transport via OATP2B1. PFOA in the intestinal venous plasma follows the portal plasma flow to the extracellular space of the liver, where it is transported to the hepatocytes via OATP1B1 and OATP1B3. From the hepatocytes, PFOA is either transported to the systemic circulation, or secreted to the bile by BSEP followed by the transport to the intestinal lumen. B) PFOA reaching the kidneys is filtrated by the glomeruli to the primary urine in the proximal tubule lumen, from where it is either excreted via the urine, or reabsorbed to the proximal tubule cells via the OAT4 transporter. From the proximal tubule cells, PFOA reaches the kidney venous plasma and thereafter systemic circulation. An overarching principle is that only the fu of PFOA passes through the different membranes, is actively transported, or filtrated by the glomeruli. We assumed an instant chemical equilibrium between the venous plasma and the extracellular space of the intestine and kidney, as well as the interstitium and the vascular space of the liver.

A representation of the model structure is given in Figure 2 and the model code is available in GitHub (Pachoulide, 2025).

**Figure 2.**
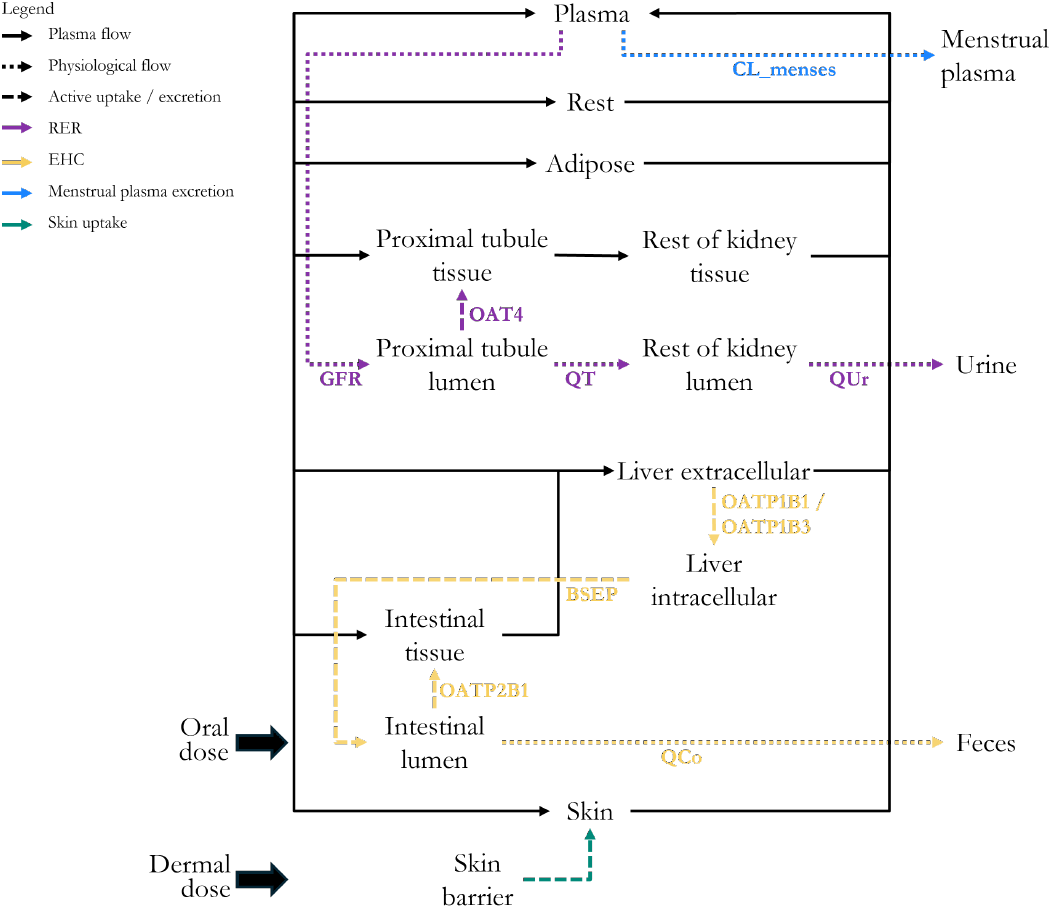
A schematic representation of the NG-PBK PBK model for PFOA for both oral and dermal exposure. Rest refers to all other tissues that were not explicitly modelled. The black solid arrows represent the plasma flow, the dotted arrows a physiological flow, and the dashed arrows represent an active process. In green is represented the skin uptake, in yellow the enterohepatic circulation, in purple the renal excretion and re-absorption and in blue the menstrual clearance.

## 4 Materials and Methods

### 4.1 General workflow

The mechanistic PBK model for simulating PFOA toxicokinetics in humans was developed with the aim of facilitating the extrapolation to other PFAS, using in vitro and in silico derived parameters. To this end, the model structure was constructed based on up-to-date knowledge of the processes driving the toxicokinetics of PFOA, as described in Section 3 and the supplementary materials. The model was subsequently translated into mathematical equations and parameterized using in vitro derived parameters for PFOA. The model was evaluated in two steps, by goodness-of-fit comparisons and by global sensitivity analysis (GSA).

### 4.2 PBK model equations

The PBK model consists of a series of mass balance differential equations simulating temporal changes in PFOA amounts in the different tissues. Detailed description of the equations in each organ can be found in the supplementary information.

The general equation for calculating the change in the amount of PFOA in an organ is 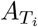 in *µg/*day, as Equation 1 and Equation 2:

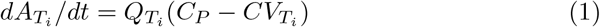

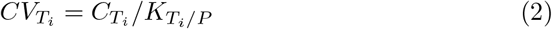

Where 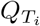 is the tissue plasma flow rate in *L/*day, *C*_*P*_ and 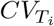 are the PFOA plasma concentration entering the tissue and that in the venous plasma leaving the tissue, respectively in *µg/L*, and 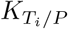 is the tissue:plasma partition coefficient.

Equation 3 was used to calculate PFOA tissue-plasma partition coefficients.

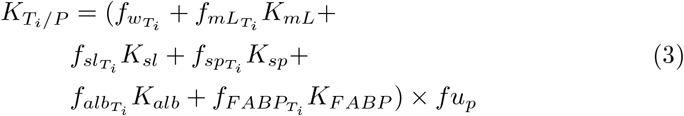

Where 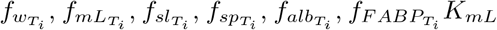, are the fractional volumes of water, membrane lipids, storage lipids, structural proteins, albumin and fatty acid binding protein in each tissue, respectively; *K*_*sl*_, *K*_*sp*_, *K*_*alb*_, and *K*_*F aBP*_ are the PFOA sorption coefficients to membrane lipids, storage lipids, structural proteins, albumin and fatty acid binding proteins; and *fu*_*P*_ is the fraction unbound of PFOA in plasma. The sorption coefficients of PFOA to the different tissue constituents were taken from (Allendorf et al., 2020) (see Theory), and the fractional volumes of the relevant tissue components were taken from (Allendorf et al., 2020; Utsey et al., 2020). The values of both the sorption coefficients and fractional volumes are available in the supplementary materials (Figures S5-7, and Table S1). A detailed description of the rationale underpinning the selection of this method for determining the partition coefficients can also be found in the supplementary materials.

The fu of PFOA in plasma was dynamically calculated depending on age, based on serum albumin concentrations, using Equation 4, previously described by (Fischer et al., 2024).

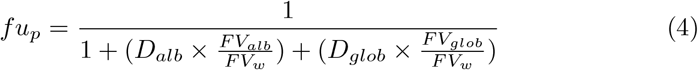

Despite that fu is one of the most critical parameters in PBK models, frequently only the fu in plasma is determined experimentally, but not that in tissues. To determine the fu in each matrix of interest, we followed the recommendations of (Poulin and Haddad, 2018), who developed an equation for calculating an adjusted fu based on the local albumin concentrations in the relevant matrices Equation 5.

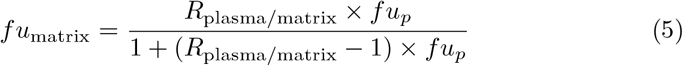

Where *fu*_matrix_ is the fraction unbound of PFOA in either liver, intestine, or proximal tubule lumen and *R*_plasma/matrix_ is the albumin concentration ratio between plasma and the matrix. The original equation by Equation 5 corrects for the local fraction unionized as well; however, given the absence of significant differences between the pH of plasma and those of the matrices of interest, it was assumed that there would be no such differences in the fraction unionized of PFOA either. Consequently, the term was removed from the equation.

PFOA uptake was modelled either from the intestine alone, or from both the intestine and the skin, using apparent permeability (Papp) values obtained in vitro (Janssen et al., 2024; Ragnarsdóttir et al., 2024), and Equation 6, where

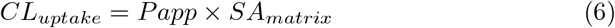

Where *CL*_*uptake*_ stands for the passive uptake clearance in *L/day, P app* stands for the apparent permeability, in *cm/s* and *SA*_matrix_ stands for the skin, or intestinal surface area, in *cm*^2^.

PFOA excretion was modelled via feces (EHC), GF and RER and clearance through menstruation. As described in Section 3, active transport determines both the EHC and RER of PFOA. The in vitro derived Vmax and transporter Km was scaled to the in vivo transport rates following in vitro to in vivo (IVIVE) extrapolation principles as suggested by (Tan et al., 2024) and equations Equation 7, Equation 8, Equation 9 :

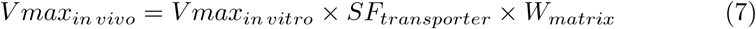

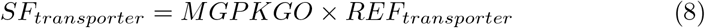

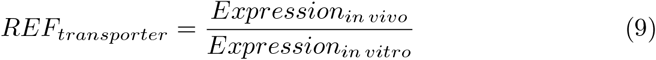

*SF*_*transporter*_ refers to the transporter specific scaling factor, *W*_*matrix*_ to the organ weight in kg, *MGPKGO* to milligram protein per kilogram organ and *REF*_*transporter*_ to the relative expression factor between the in vitro system and the in vivo situation. *OATP1B1, OATP1B3, OATP2B1 and OAT4* transporter expression in the in vitro systems and the relevant organs were taken from literature (Lin et al., 2023; Al-Majdoub et al., 2021). A consolidated description of the parameter scaling, including specific references and assumptions is found in Table 1.

**Table 1.** Parameter scaling for transporter activities. ^1^,^2^: OATP2B1 and OATP1B1/1B3 (Lin et al., 2023), BSEP assumed the same as TCA (de Bruijn et al., 2024), OAT4 (Louisse et al., 2024) ^3^: milligram protein per kilogram intestine, liver and kidney respectively (Utsey et al., 2020), mg protein per 10*e*^6^ hepatocytes and number of hepatocytes per kilogram liver (de Bruijn et al., 2024) ^4^: calculated using equation 9; mg protein/mg membrane in vivo / mg protein/mg membrane in vitro; OATP2B1: 1.6 (assuming the same as the liver)/0.812 (Lin et al., 2023), OATP1B1: 2/0.120 (Lin et al., 2023), OATP1B3: 1/0.719 (Lin et al., 2023), BSEP: assumed as 1, OAT4: 1.9 (Al-Majdoub et al., 2021)/0.063 (Lin et al., 2023) ^5^: calculated using equation 8 ^6^: scaled to the weight of the intestinal lumen, liver extracellular space, liver intracellular space and proximal tubule lumen respectively; in this example, the fractional volumes and a body weight were taken from table S2 ^7^: calculated using equation 7, MW = 414.07g/mol

|  | In vitro |  | Scaling |  |  |  | In vivo |  |
| --- | --- | --- | --- | --- | --- | --- | --- | --- |
|  | Vmax<sup>1</sup><br>(umol/min/<br>mg protein) | Km<sup>2</sup><br>(umol/L) | MGP KGO<sup>3</sup><br>(g protein/<br>kg organ) | REF<sup>4</sup> | SF<sup>5</sup> | KGO<sup>6</sup><br>(kg<br>organ) | Vmax<sup>7</sup><br>(ug/day) | Km<br>(ug/L) |
| Intestine<br>(OATP2B1) | 0.0015 | 148.68 | 1.3e5 | 1.97 | 2.56e5 | 3.08 | 7.05e8 | 6.16e4 |
| Liver<br>(OATP1B1) | 0.0023 | 52.65 | 1.8e5 | 16.67 | 3.00e6 | 0.84 | 3.46e9 | 2.18e4 |
| Liver<br>(OATP1B3) | 0.0027 | 91.61 | 1.8e5 | 1.39 | 2.50e5 | 0.84 | 3.38e8 | 3.79e4 |
| Liver<br>(BSEP) | 7 | 16.4 | 11.62 | 1 | 11.62 | 1.13 | 5.48e7 | 6.79e4 |
| Kidney<br>(OAT4) | 4.5e-3 | 47 | 1.7e5 | 30.16 | 5.13e6 | 0.13 | 1.789e9 | 1.954e4 |

Combining the albumin mediated active transport and the *Michaelis-Menten* equation, active transport was calculated as shown in equation Equation 10 :

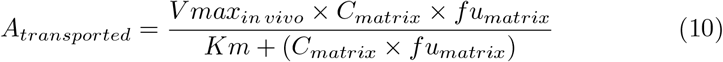

*A*_*transported*_ is the amount of PFOA that is transported at a given time-point, *C*_*matrix*_ is the concentration of PFOA in the matrix where it is transported from and *fu*_*matrix*_ is the fraction unbound of PFOA at the matrix where it is transported from.

GFR was estimated using the full age spectrum (FAS) equation (Pottel et al., 2016) (Figure S2) Equation 11 :

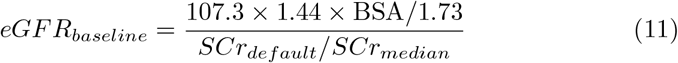

*eGF R*_*baseline*_ stands for the baseline GFR, *BSA* stands for the body surface area, and *SCr*_*default*_ and *SCr*_*median*_ stand for the default and median serum creatinine (SCr) values. Default SCr values for adults were 0.90 and 0.70 for males and females, respectively. Median GFR values per age and sex were estimated using Equation 12 and the age-specific GFR was estimated using Equation 13, which describes an exponential decrease in GFR after 40 years of age.

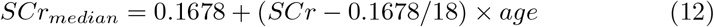

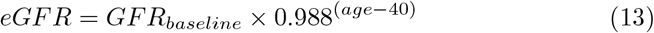

Plasma clearance via menstrual blood was modelled as previously done by (Ruark et al., 2017). Given that total plasma is eliminated during menstruation (Ruark et al., 2017), it was assumed that: i) PFOA concentrations in plasma and menstrual excretion fluid were equal, and ii) total PFOA plasma concentration (and not the fu) was cleared via this route. Menarche and menopause were modelled to start at 12 and 51 years of age, respectively. Plasma clearance was at 25 mL per cycle for women of 12-to 45-years of age and then at 27.1 mL per cycle in women from 45-to 51-year-old women. Cycle lengths were of 29.2 days from age 12-to 45-years and then of 35.1 days, until menopause.

### 4.3 PBK model parameters

Tables S2 and S3 compile all model parameters for an adult woman, their values, units and references. Organ volumes and flow rates were calculated based on the fractional organ volumes and flow rates, plasma cardiac output and body weight. Equations to calculate the aforementioned parameters for both sexes over a lifetime were taken from (Ratier et al., 2024), except for the equations to calculate body weight, body height, and body surface area which were taken from (Gastellu et al., 2025). The fractional volume of the intestinal lumen was calculated from (Willmann et al., 2004), the fractional volumes of the proximal tubule tissue, proximal tubule lumen, rest of kidney tissue and rest of kidney lumen were derived from (Pletz et al., 2021). Fractional volumes of liver extracellular and intracellular space were derived from (Utsey et al., 2020). Serum albumin concentration was modelled to dynamically change during lifetime. For this we derived a polynomial equation, based on average values from (Weaving et al., 2015) (see Equations S1 and S2 and, Figure S1). The tubular fluid flow and urinary excretion rate were taken from (Scotcher et al., 2016; Pletz et al., 2021). Compound-specific parameters such as sorption coefficients to tissue components, plasma albumin and globulins, apparent permeabilities across the skin and intestine, and Vmax and Km were taken from (Allendorf et al., 2020; Fischer et al., 2024; Ragnarsdóttir et al., 2024; Janssen et al., 2024; Lin et al., 2023; de Bruijn et al., 2024; Louisse et al., 2024).

### 4.4 Model Evaluation

The NG-PBK model was evaluated following the recommendations of the OECD PBK guidance document (OECD, 2021a), which recommends to perform a sensitivity analysis to quantify the extent at which the uncertainty of the model can be attributed to the input parameters, and an evaluation of the goodness-of-fit where the PBK model predictions were compared to observed in vivo kinetic data (OECD, 2021a).

#### 4.4.1 Goodness-of-fit

We evaluated the predictions of the developed NG-PBK model against: a) predictions by previously published PBK models (EFSA, 2020; Ratier et al., 2024), b) a human volunteer study (Abraham et al., 2024) and c) human biomonitoring data (Olsen et al., 2007; Johanson et al., 2023; Bartell et al., 2010; Seals et al., 2011; Worley et al., 2017; Xu et al., 2020; Zhang et al., 2013; Li et al., 2017; Batzella et al., 2024; Fu et al., 2016). The model outputs used for the evaluation step were individual serum concentrations, cumulative PFOA amount in urine and faeces and/or predicted half-life. We evaluated the goodness-of-fits either visually and/ or quantitatively via a linear regression analysis. A two-fold difference between model estimates and observed data, or a difference within the intrinsic variability of the measured data were considered acceptable. It is noted that given that PFOA serum to plasma ratio is 1, plasma and serum concentrations were assumed equal, and the terms were used interchangeably.

##### 4.4.1.1 Previously published PBK models

The predictions of our model were compared to those of the model optimized by EFSA, “EFSA-2020” model (EFSA, 2020). The two models were run with the same input settings, corresponding to the (Abraham et al., 2024) study. The model predictions were also evaluated against those of the model published by (Ratier et al., 2024), “Ratier-2024” model, which simulates PFOA toxicokinetics in pregnant women, infants, children, and adolescents. This enabled us to evaluate the influence of the updated physiological parameters and model structure on model predictions, as well as the computational efficiency of the model. To this end, the two models were run with the predicted daily intakes of each subject of the HELIX cohort, obtained in (Ratier et al., 2024) via reverse dosimetry in a Bayesian framework. The HELIX cohort consisted of 1,239 children from 6 European countries (France, Greece, Spain, Lithuania, Norway, and the United Kingdom). All subjects were simulated with an exposure to PFOA for 1 day, which was followed by a wash-out period of 30 days.

##### 4.4.1.2 Human volunteer study

One controlled human study was available in literature which aimed to determine the toxicokinetic profile of PFAS, including PFOA, after oral exposure (Abraham et al., 2024). The study subject was a 67-year-old male, who ingested a mixture of radiolabeled PFOA. PFOA serum, urine and faeces samplings were taken over 450 days at multiple time points (Abraham et al., 2024). We simulated a male subject with the same physiological characteristics as the subject of the human study (67 years old, with a body weight of 82 kg), being exposed to a single oral PFOA exposure of 0.048 *µ*g/kg bw.

##### 4.4.1.3 Human Biomonitoring studies

Lastly, we compared the model predicted serum concentrations to those measured in HBM studies (Johanson et al., 2023; Olsen et al., 2007). The studies involved a total of 131 subjects that were exposed to PFOA either via drinking water (Johanson et al., 2023), or occupational exposure (Olsen et al., 2007). We compared the predicted PFOA serum concentrations with those measured, and the predicted half-lives with those reported in literature (Olsen et al., 2007; Johanson et al., 2023; Bartell et al., 2010; Seals et al., 2011; Worley et al., 2017; Xu et al., 2020; Zhang et al., 2013; Li et al., 2017; Batzella et al., 2024; Fu et al., 2016). Given that the inter-individual variability of the measured concentrations was about ten-fold, we considered that a ten-fold difference between measured and estimated datasets was acceptable. In all studies, PFOA serum concentrations were measured after PFOA exposure ceased. For the simulations, physiological information of each subject was estimated based on their reported sex and age. Exposure and simulation time were subject-specific and were determined based on the reconstructed exposure estimations.

As the exact exposure was unknown, this was reconstructed following two different approaches. For the subjects in the (Johanson et al., 2023) study (“Arvidsjaur”, “Lulnäset” and “Visby”) PFOA exposure was assumed to be a combination of contaminated drinking water and background food intake. For the population in the (Olsen et al., 2007) study (“Olsen”), the exposure route was not available, but PFOA serum concentrations were available from two points in time (at the point of retirement and about five years later). Based on the available information, we used the measured PFOA serum concentration at the first measurement as a starting point to estimate the exposure dose by reverse dosimetry; and the second measurement as the observed serum concentration against which the model predicted serum concentration was evaluated.

#### 4.4.2 Global Sensitivity Analysis (GSA)

To perform GSA, we followed the workflow first introduced by (McNally et al., 2011) and then adopted by (OECD, 2021a). The workflow consisted of a two-step sensitivity analysis: starting with an elementary effect screening test (Morris Test) followed by a variance-based test (eFAST). The Morris Test was performed with PFOA plasma concentration as the model-output, and the eFAST with PFOA plasma, proximal tubule tissue, proximal tubule lumen and liver intracellular concentrations as the model-outputs. Given the long half-life of PFOA, a 20-year simulation was chosen to allow for the full elimination of PFOA from the body after a single oral dose. This assumption is based on a half-life of 4 years for PFOA and reaching an apparent steady-state approximately after 5 half-lives. To enable convergence of the outcomes, the experimental designs consisted of 1,000 elementary effects per parameter for the Morris test, and 10,000 runs per retained parameter in the eFAST test. This led to 2,924 and 360,000 simulations of the PBK model, respectively. The test designs, parameter distribution and script are provided in GitHub (Pachoulide, 2025).

The Morris elementary effect test categorized the parameters depending on their sensitivity indices: mean effect (*µ*^∗^) and standard deviation (*σ*). The values of the sensitivity indices were relative, meaning that the sensitivity index of a parameter was high relative to that of other parameters. All parameters with relatively small (*µ*^∗^) and (*σ*) were considered non-influential to the model’s output. The parameters with a global elementary effect above 5% were applied to the eFAST, which further ranked the parameters based on their total order index. Total order index represented the total variance of the output caused by both the variance of a specific parameter (main effect) and that parameter paired with another (interaction) (Saltelli et al., 1999). The final model had a total number of 52 parameters, and all of them were included in the Morris Test. Parameters were assumed to follow a uniform distribution, and varied by 10% in both directions, for both tests. Parameter baseline default and probability distribution values can be found in GitHub (Pachoulide, 2025).

Lastly, we performed a Pearson correlation analysis to study the relationship between the predicted half-life and different physiological parameters that vary over lifetime. These parameters were: the age at exposure (expAGE), body weight (BW), GFR, fu in plasma (fup) (which depends on the differences in serum albumin concentration over lifetime), and PFOA clearance through menstruation (CL_menses). For this analysis, a group of 16 males and females was created, with ages at the beginning of exposure varying from 0-to 64-years. All simulated subjects were exposed to 0.00019 *µ*g/kg BW/day of PFOA for one year, which was followed up by a wash-out period of 15 years.

### 4.5 Codes and software

All model and data analysis codes are available in GitHub (Pachoulide, 2025). The PBK model was written in the R language and run using RStudio, version R 4. 5. 1. The deSolve_1.40, PKNCA_0.12.1, pracma_2.4.4 and sensitivity_1.30.2 packages of R were used for running the model, calculating the half-life, AUC and performing the sensitivity analysis, respectively. Data structuring, manipulation and plotting were performed within the tidyverse_2.0.0 package of R as well.

## 5 Results and Discussion

The developed NG-PBK model for PFOA is the first PFAS toxicokinetic model that integrated various parameters derived from in vitro experiments and in silico approaches. An illustration of predicted PFOA internal concentrations over lifetime, in both sexes can be found in SI Figures 9-11. We demonstrated the model’s predictability against HBM data covering various life-stages and exposure ranges and identified influential parameters by global sensitivity analysis. The evaluation of the model’s performance against HBM data is one of the practices recommended within the NGRA framework.

### 5.1 Model evaluation

#### 5.1.1 Comparison against reference models

The NG-PBK model’s predictive capacity was evaluated by means of comparison to previously published and evaluated PFOA PBK models, here referred to as reference models. The two reference models were the model used in the risk assessment process, “EFSA-2020” (EFSA, 2020) and that of (Ratier et al., 2024), which predicted PFOA concentrations throughout the different life-stages and especially infants and children. As shown in Figure 3 and SI Figure 12, the NG-PBK model predicted the PFOA plasma concentration similarly to both reference models, with the majority of predictions falling within the recommended two-fold difference range (WHO, 2010). Furthermore, while the predicted liver and kidney concentrations between the two models were different (Figure S12), in the absence of data pairing tissue and plasma concentrations of PFOA in humans at steady state, it is challenging to evaluate the accuracy of the predicted PFOA tissue concentrations. Nevertheless, at predicted plasma concentrations corresponding to those measured in human autopsies by (Nielsen et al., 2023), both the kidney and liver concentrations predicted by the EFSA-2020 model were overestimated. On the contrary, the NG-PBK model seemed to only mis(under)-estimate the liver concentrations (Figure S13). The discrepancy between the predicted and observed liver concentrations could be attributed to limitations of both the reference dataset of human autopsies, or the model.

**Figure 3.**
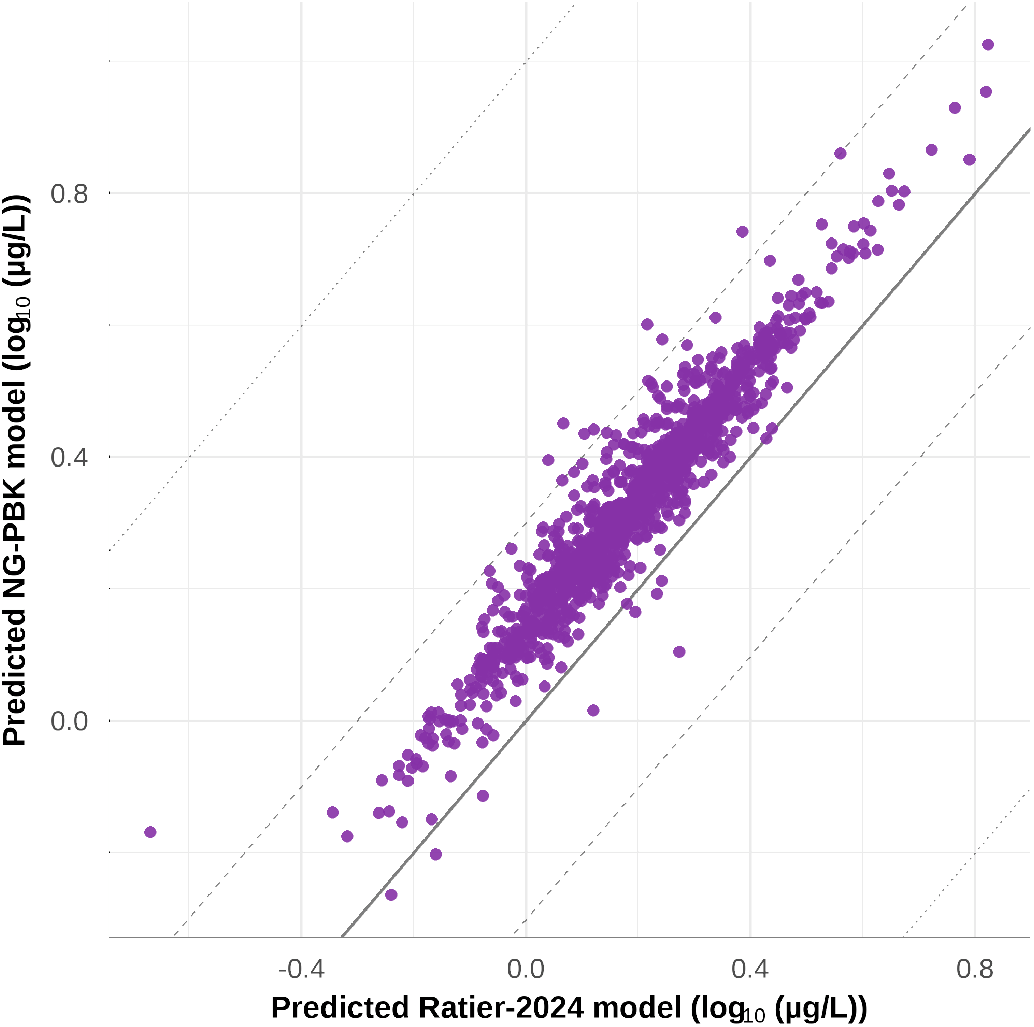
Evaluation of the NG-PBK model against the reference Ratier-2024 model (Ratier et al., 2024). Predicted PFOA plasma concentration from the Ratier-2024 PBK model is plotted against that predicted by the NG-PBK model. Solid, dashed, and dotted lines represent perfect alignment, two-fold, and ten-fold difference, respectively.

Figure 3 demonstrates that predicted PFOA plasma concentrations from both models follow the same trend. Although plasma concentrations predicted by the NG-PBK model were, on average, 1.39 times higher than those predicted by the Ratier-2024 model, this discrepancy is within the acceptable two-fold range. Furthermore, even though a few points are outside the two-fold range difference, the maximum discrepancy between the two sets of predictions is 3.15-fold Figure 3. The slight over-prediction of the NG-PBK model can be attributed to small differences in the calculation of the physiological parameters over lifetime, the PBK-model structure and input parameters. Overall, these results demonstrated comparable performances of the NG-PBK model to the two reference models which were calibrated on in vivo toxicokinetic data.

#### 5.1.3 Comparison against a human controlled toxicokinetic study

Next, we compared PFOA plasma concentration measurements from a human-controlled toxicokinetic study with the predictions by the NG-PBK model and EFSA-2020 model. As shown in Figure 4, the predicted PFOA plasma concentrations were close to those measured in (Abraham et al., 2024). The NG-PBK model predicted the measured plasma concentrations slightly better than the the EFSA-2020 model. However, the predicted maximum concentration (Cmax) in plasma, from both models (0.38 and 0.36 ng/mL for the EFSA and NG-PBK models, respectively), was underestimated compared to the observed one (0.8 ng/mL). Both models appeared to miss the initial high Cmax and slow tissue distribution observed in the human study (Abraham et al., 2024). Nevertheless, we believe that this discrepancy should not influence the adequacy of the model for its application in the risk assessment process. Due to the facts that chronic (and not acute) exposure is considered critical for the risk assessment of PFOA (EFSA, 2020), and that oral bioavailability is highly dependent on the food matrix upon intake, nutritional habits, microbiota composition, metabolism, age, sex and BMI (Heitmann et al., 2021; Fischer and Fadda, 2016).

**Figure 4.**
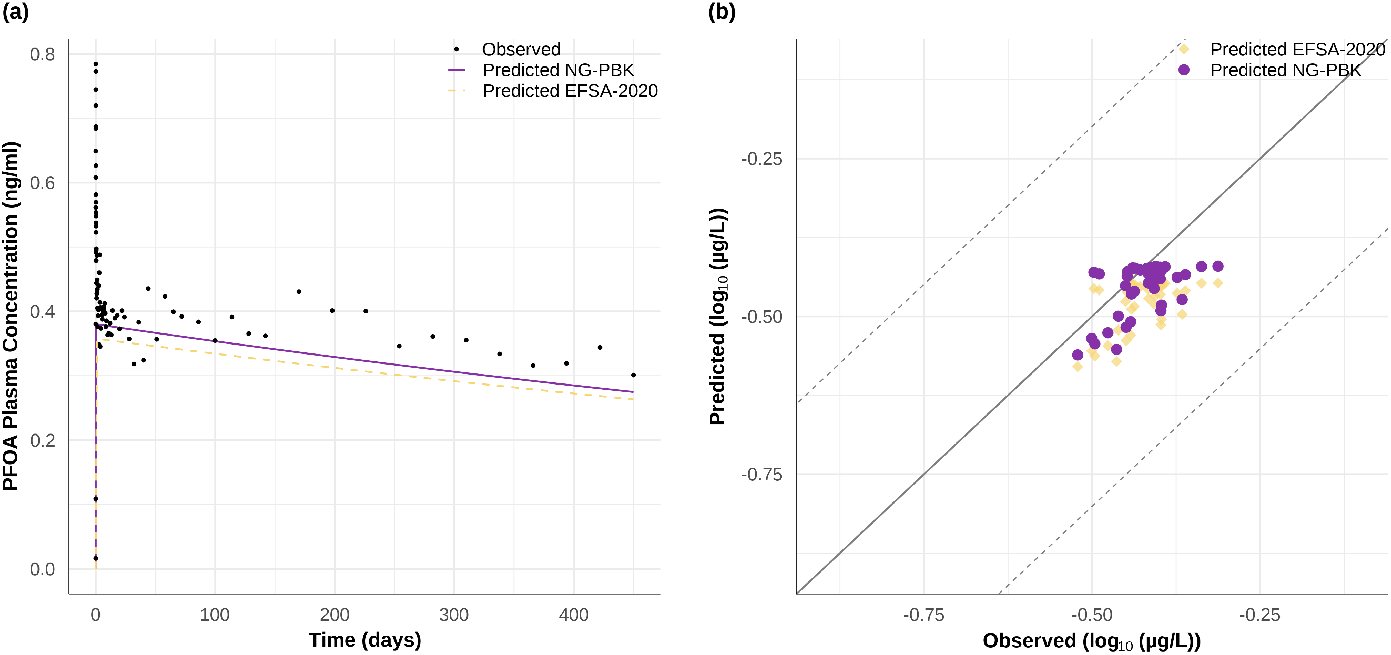
Evaluation of the NG-PBK model against the human volunteer study (Abraham et al., 2024) and the reference EFSA-2020 model (EFSA, 2020). Panel (a) compares the observed and modelled plasma concentration of PFOA over time. The observed (dark points), predicted by the NG-PBK model (solid purple line) and predicted by the EFSA-2020 model (yellow dashed line) plasma concentrations are plotted in overlay. Panel (b) compares observed and NG-PBK model (purple points), or EFSA-2020 model (yellow diamonds) predicted PFOA plasma concentrations. Solid and dashed lines represent perfect alignment and two-fold difference, respectively.

#### 5.1.3 Comparison against human biomonitoring studies with uncertain exposure information

We further evaluated the capacity of the NG-PBK model to predict PFOA serum concentrations measured in four HBM studies : Arvidsjaur, Lulnäset and Visby published by (Johanson et al., 2023), and Olsen initially published by (Olsen et al., 2007). Detailed description of the re-construction of the exposure estimates for each study can be found in the supplementary materials (Figure S15). For the Arvidsjaur, Lulnäset and Visby subjects, the median estimated oral exposure was at 0.0007 *µ*g/day/kg BW ranging from 0.0002 to 0.0075 *µ*g/day/kg BW. For the Olsen subjects, the median estimated dermal exposure was at 0.0310 *µ*g/day/kg BW, with a range from 0.0055 to 0.3975 *µ*g/day/kg BW.

As depicted in Figure 5, panel (a), the predicted PFOA concentrations were in general slightly higher than the observed ones. For the majority (95%) of the data points, the discrepancy between the observed and predicted PFOA plasma concentration was less than 10-fold. A 10-fold range was considered acceptable in this context. For half of the data points, the discrepancy between the observed and predicted plasma concentrations falls within the 2-fold range recommended by the WHO (WHO, 2010). It should also be noted that there are 7 subjects for which the predicted and observed plasma concentrations have more than 10-fold difference. However, these subjects are considered outliers, because their serum concentrations were either extremely high or low, and did not correlate with their estimated exposure.

**Figure 5.**
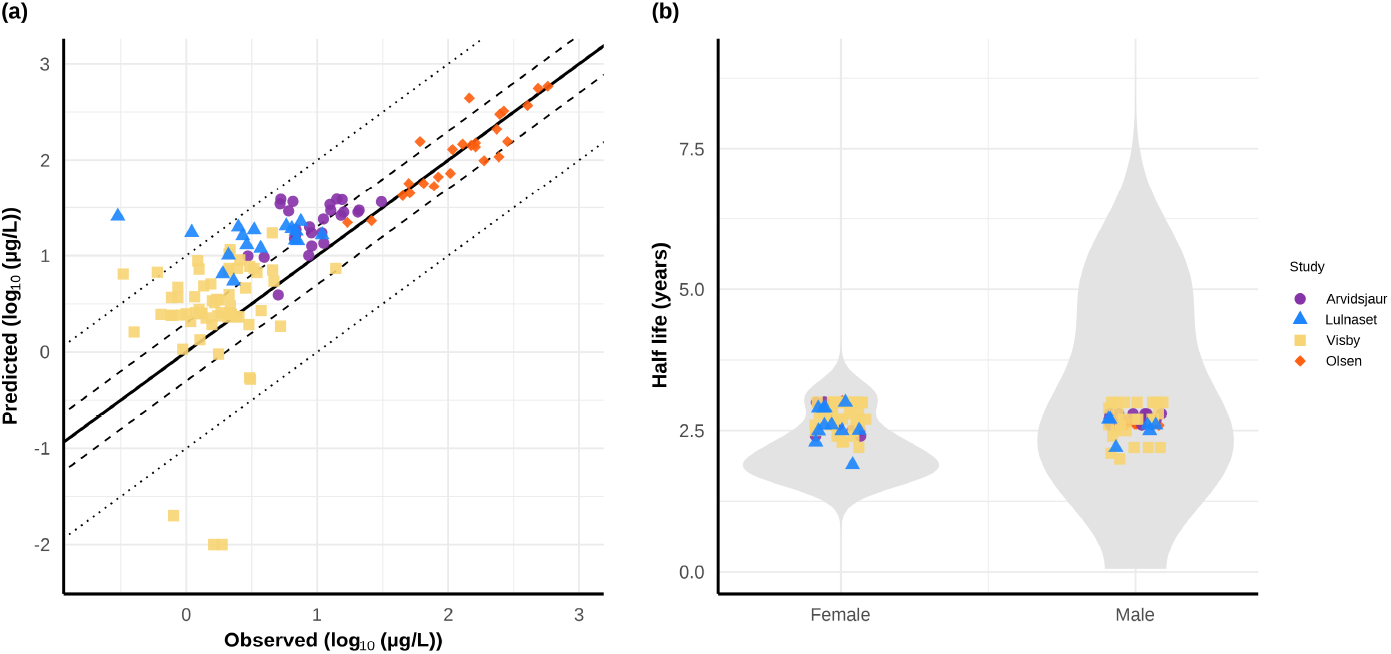
Integration of the NG-PBK model with HBM based exposure data. Panel (a) compares observed and NG-PBK model predicted PFOA serum concentrations. Solid, dashed, and dotted lines represent perfect alignment, two- and ten-fold difference, respectively. Panel (b) illustrates the range of reported PFOA half-lives taken from literature over which are shown the predicted half-lives. Purple circles, blue triangles, yellow squares, and green diamonds represent subjects from the Arvidsjaur, Lulnäset and Olsen studies, respectively.

We further evaluated the distributions of predicted PFOA half-lives against those reported in literature for each sex (6250 data points for men, 6360 data points for women) as violin plots, shown in panel (b) of Figure 5. For the literature-reported data, we observed that there is a more widespread distribution of reported half-lives in men compared to women. This highlighted the high inter-individual variability in reported half-lives. For both sexes, two peaks of high distribution density were identified, around 2 or 3 years for women, and 2.5 or 5 years for men. Overall, it appears that women have shorter half-lives than men.

On top of the violin plots are superposed data points of the predicted half-lives from the evaluated HBM studies, placed well within the range of previously reported half-lives, and around the values with the highest density Figure 5. Interestingly, most of the predicted half-lives are around 2 to 3 years, irrespective of the sex and the inter-individual variability is narrower in the predicted half-lives compared to those reported in literature. This could be attributed to unknown background exposure to PFOA, or inter-individual differences in elimination capacity (such as physio-pathological conditions affecting the kidney, liver, serum albumin, or lipid to protein ratio), for which the model and input data do not account for. Overall, these results demonstrate the model’s capacity to accurately predict PFOA plasma concentrations based on reconstructed exposures for each individual.

### 5.2 Global Sensitivity Analysis

We evaluated the model following the OECD guidance on the characterization, validation, and reporting of PBK models for regulatory purposes (OECD, 2021a). The guidance recommends a two-step GSA, firstly an elementary effect screening test (Morris Test) and secondly a variance-based test (eFAST). For the Morris test, PFOA concentration in plasma was the model output for the test, while for eFAST the concentration of PFOA in plasma, proximal tubule tissue, proximal tubule lumen and liver intracellular space were taken as evaluation outputs. Results are shown in Figure 6 and SI Figures 16-17.

**Figure 6.**
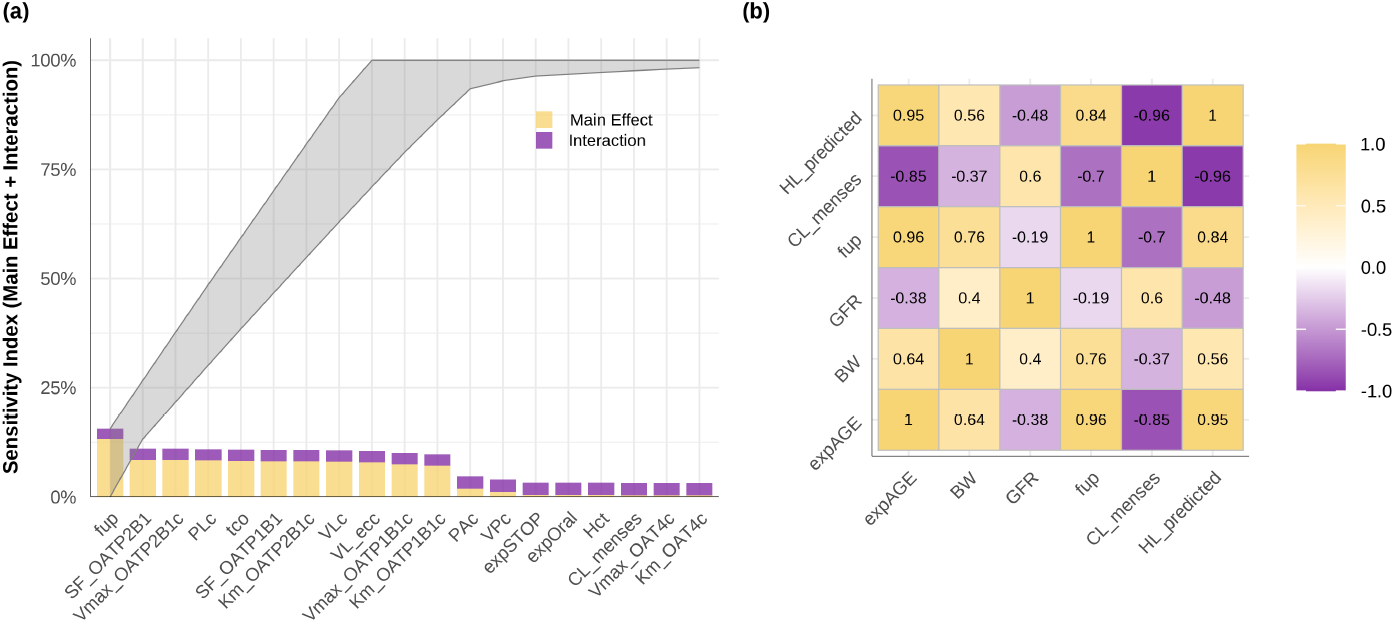
Results of the global sensitivity and Pearson correlation analysis. Panel (a) shows the Lowry plot showing the results of the eFAST analysis. Sensitivity indices presented here were calculated accounting for the sensitivity of the predicted PFOA plasma concentration to the input parameters. A simulation time of 20 years, with steps of one month was performed, covering all kinetic processes (absorption, distribution, and elimination). The main effect (yellow) and interaction (purple) are stacked on top of each other, illustrating the total effect of each parameter. The grey ribbon represents the cumulative effect of all parameters on the model output. Panel (b) shows the correlation plot between physiological parameters (age at exposure (expAGE), body weight (BW), GFR, fu in plasma (fup) (dependent on the differences in serum albumin concentration over lifetime), and PFOA clearance through menstruation (CL_menses)) with the predicted half-life of PFOA (HL_predicted) in females. The higher the correlation the darker the colour becomes, with the yellow colour illustrating a positive correlation and the purple colour a negative correlation.

Results from the Morris test suggested that PFOA plasma concentration was primarily sensitive to the fraction unbound in plasma (fup), parameters influencing the enterohepatic circulation and to a lesser extent to parameters influencing renal excretion and reabsorption (Figure S16). To further explore the effect of each parameter, the parameters with an elementary effect score above 5% were also evaluated using the eFAST variance test. The eFAST test was run on 36 parameters and convergence was only reached with a sample size of 10,000 (32 hours of simulation time). Results of the eFAST are illustrated on the Lowry plot, Figure 6, panel (a).

As shown in Figure 6 and SI Figure 17, all of the evaluated model outputs were sensitive to the same parameters. These were the fraction unbound in plasma (fup) and parameters related to the enterohepatic circulation (i.e. SF_OATP2B1, Vmax_OATP2B1 and Km_OATP2B1 driving the intestinal uptake; colon transit time driving the faecal excretion, and SF_OATP1B1, the volume of liver and liver:plasma partition coefficient which influence the active uptake and elimination from the hepatocytes). Although the most sensitive parameters were consistent for all model outputs, their sensitivity index and ranking differed for each model output. Interestingly, the ranking of these parameters was identical when the concentration of PFOA in plasma and proximal tubule were the evaluated model outputs, with the fup being the most highly sensitive parameter. On the contrary, the ranking was different when liver intracellular concentration was selected, with the fup being ranked as the sixth most sensitive parameter.

To further evaluate the processes determining the model predictions, we also studied the correlation between the predicted half-life (one of the most discussed toxicokinetic characteristics of PFOA) and input parameters related to physiological changes during lifetime. The results shown in Figure 6 and SI Figure 18, demonstrate that CL_menses, expAGE and fup had high correlation scores (-0.96, 0.95, and 0.84, respectively), while BW and GFR had lower scores (0.56 and -0.48). These results are indicative of the influence of menstruation and age on the half-life of PFOA, as previously demonstrated in epidemiological studies (Rosato et al., 2024; Batzella et al., 2024). The positive relationship between fraction unbound in plasma and half-life is unexpected, as one would expect that elimination is increased, and therefore half-life decreased with an increased fraction unbound. However, an increase in the fup could also lead to an increase of PFOA amounts in tissues, and a decrease in the amount of PFOA lost via menstruation. This is also reflected in the results of the eFAST analysis where the fraction unbound in plasma was also a parameter highly interacting with other input parameters.

Both the GSA and parameter correlation results suggested that the rate-limiting parameter in the RER processes was the fu in plasma, which was supported by the following three observations. Firstly, the fraction unbound in plasma was consistently the parameter to which PFOA concentrations in plasma, the proximal tubule tissue, and the proximal tubule lumen were the most sensitive to (Figure S16). Secondly, the sensitivity indices of both GFR and OAT4 Vmax and Km were not among the most highly ranked, in comparison to fu. Thirdly, the correlation analysis demonstrated an identical correlation of both parameters to the predicted half-life Figure 6. Contrary to our results, renal reabsorption parameters (Kt, Tmax), and PFOA half-life were two of the most sensitive parameters in previous PBK models (EFSA, 2020; Loccisano et al., 2011; Husøy et al., 2023; Lin et al., 2023; Ratier et al., 2024). This can be explained by the fact that the renal reabsorption parameters in the previously published PBK models were fitted to observed half-lives, potentially biasing the model towards this parameter (EFSA, 2020; Loccisano et al., 2011; Husøy et al., 2023; Lin et al., 2023; Ratier et al., 2024).

This is the first PBK study that showed that the influence of enterohepatic circulation on the predicted plasma concentrations was higher than that of renal elimination and reabsorption in humans. These results are aligned well with the mechanistic PBK model of PFAS in mice that showed that enterohepatic circulation was driving the plasma concentration of PFAS with a chain length with more than 7 carbons (Fischer et al., 2025). Similarly, the higher influence of enterohepatic circulation and faecal elimination compared to renal clearance, in long chain PFAS, including PFOA, was also observed in human volunteer studies (Abraham et al., 2024; Fujii et al., 2015).

Our modelling results were also in agreement with epidemiological studies that suggested that PFOA clearance via menstruation was one of the contributing factors explaining the shorter half-lives and lower measured plasma concentrations in women compared to men. As seen in Figures S10 and S11, our model correctly captured the observed lower plasma concentrations in women compared to men, especially during the menstrual years of women (12-51 years of age). These results coincided with previous modelling work, where simulated PFOA serum concentrations were higher in post-menopausal women compared to pre-menopausal ones (Ruark et al., 2017).

### 5.3 Limitations and future developments

The present study introduces a NG-PBK model that relies exclusively on NAM-derived input parameters and has not been calibrated to human data. Nevertheless, it is important to acknowledge that this approach is also the source of the model’s limitations. The NG-PBK model-predicted plasma concentrations were slightly overestimated against those measured in the human volunteer and HBM studies (Abraham et al., 2024; Olsen et al., 2007; Johanson et al., 2023), and the predicted liver concentrations, were lower than those measured in human autopsy samples by (Nielsen et al., 2023), (Figure S13). In the subsequent section, we discuss a range of potential explanations for these discrepancies.

Measured PFOA plasma concentrations in HBM studies were systematically over-predicted by the NG-PBK model. The reasons for this result might largely relate to the reconstruction of exposure estimations. Some limitations in the exposure estimations might be related to uncertainties about historically background exposure from food or other exposure sources over the exposure duration, potential fluctuations in PFOA concentrations in drinking water, inter-individual variability in drinking water consumption, as well as dynamic individual behaviours (i.e., movements between different residencies). Moreover, the high inter-individual variability in the observed half-lives, but also serum concentration could be partly explained by physio-pathological conditions, or inhibition/induction of the active transporters, which are not taken into consideration in the NG-PBK model.

Physio-pathological conditions that could explain the observed high interindividual variability are conditions that alter the serum albumin concentrations (such as age, pregnancy, or obesity), or conditions altering hepatic or renal function (such as diabetes, chronic kidney disease, metabolic syndrome, or cholestasis). An increasing amount of evidence shows an association between exposure to PFAS, chronic kidney disease (CKD) and type 2 diabetes (Li et al., 2025; Lin et al., 2021, 2024). The studies focussing on this association showed that not only do PFAS seem to increase the risk of developing these diseases, but their toxicokinetic profiles can be also influenced by these diseases (for example a higher PFAS concentrations in CKD patients with lower GFR).

Other conditions that could be influencing PFOA biodistribution can be related to concomitant exposure to xenobiotics such as medicines, foods, smoking, and alcohol consumption. For instance, smoking and alcohol consumption were highly associated with the extend of PFOA serum elimination in humans (Batzella et al., 2024). This association is likely mediated via a decrease in the expression, or activity OATP or OAT transporters, an effect that has been demonstrated both in vitro and in vivo. Some examples are the inhibition of the expression and activity of OATP and OAT4 transporters in HEK cells, and the increased risk of developing resistance to imatinib, a drug transported by OATPs, by cigarette smoke (Sayyed et al., 2016, 2017; Mohammadi et al., 2021).

The discrepancy between the predicted and observed liver concentrations could be attributed to limitations of both the reference dataset of human autopsies, or the PBK-model. Tissue distribution of PFOA can differ between sexes and ages due to changes in organ blood flow, volumes and compositions, especially at the young or elderly ages (Li and Xiong, 2023). This physiological variability could have contributed to the observed underestimation of liver concentrations, especially because the human autopsies originated from adults of 29 to 84 years of both sexes, while the simulated subject was a single 50 years old man. Other contributing factors could be the small size of the autopsy samples (potentially not representative of whole tissue concentrations), and post-mortem differences in tissue perfusion. PBK-model-related factors could be a potential under-estimation of the oral bioavailability, and/or over-estimation of the faecal excretion.

The NG-PBK model predicted higher faecal clearance (0.078 mL/day/kg) than that estimated from human data (0.0007, 0.022 or 0.052 mL/day/kg) (Abraham et al., 2024; Andersson et al., 2025; Fujii et al., 2015) (see Table S4 and Figure S14). Nevertheless, the discrepancy was within the variability between the fecal clearance rates estimated from human studies. The discrepancies in the estimated faecal clearance may be explained by the different matrices in which PFOA was measured (i.e., fresh/freeze-dried faeces, or bile), the differences in analytical detection limits and the different methods applied for estimating faecal clearance. We believe that harmonizing these methods will improve their robustness and utilization in NGRA.

In all studies, faecal clearance was estimated assuming a complete EHC, therefore a fraction reabsorbed close to 1 (Abraham et al., 2024; Andersson et al., 2025; Fujii et al., 2015). Extensive EHC of PFOA was also supported in the EFSA risk assessment of PFAS, and was suggested to be mediated via the bile-acid transporters: *ASBT, OST, NTCP* and *BSEP* (EFSA, 2020; Cao et al., 2022), SI Figure 8. We based our choice of the parameters used to describe EHC in the NG-PBK model on the current knowledge on the expression and activity of these transporters in the human body, and available active transport kinetic values (supplementary materials, section 1.5). We assumed that *OATP2B1*, and not *OST*, contributes to the intestinal uptake of PFOA, while *OATP1B1* and *OATP1B3*, and not *NTCP*, contribute to its liver uptake. As no mechanistic data on the biliary secretion of PFOA was available, we assumed equal excretion of PFOA by the *BSEP* transporter as that of the endogenous bile acids (de Bruijn et al., 2024). Nevertheless, these assumptions should be re-evaluated upon the availability of more expression or kinetic data.

Notably, we considered that contribution of OATP2B1 to the intestinal uptake of PFOA ambiguous. While both the protein and gene expression of OATP2B1 have been detected in the intestinal tissue, there is yet no scientific consensus regarding its localization (apical, or basolateral) and activity (Oswald, 2019; Kinzi et al., 2021; Müller et al., 2017). Furthermore, given the high affinity of PFOA to the membrane phospholipid bilayer (Ebert et al., 2020), it can be expected that passive permeability contributes to a higher extent to the intestinal uptake of PFOA compared to what is currently predicted by the NG-PBK model. This underprediction could be attributed to an underestimation of the apparent permeability (Papp), due to limitations inherent in the in vitro system applied to derive the value, but also the IVIVE approach of our study.

We applied the Papp value derived by (Janssen et al., 2024) using the Epiintestinal microtissues, representing the most physiologically relevant in vitro system and the highest PFOA Papp value available. However, the in vitro system was not able to exhibit the aqueous boundary layer (ABL), which is hypothesized to enhance the uptake of ionizable chemicals like PFOA and other PFAS (Sugano et al., 2010; Ebert et al., 2024), hence leading to an underestimation of the Papp value. According to (Ebert et al., 2024) and (Dahley et al., 2023), PFAS, could enhance the permeability of the intestinal membrane by altering the pH gradient of the ABL. We therefore suggest that for future PFAS in vitro apparent permeability determination experiments, researchers implement the revised iso-pH method developed by (Dahley et al., 2023), which would allow to further explore the extent of the influence of the ABL.

According to the GSA the scaling factors (SFs) used to extrapolate in vitro transporter kinetic parameters to in vivo ones were among the most sensitive parameters of the model. Although, these SFs are uncertain due to the lack of data on transporter expression and activity in humans and in vitro systems. Importantly, neither the expression nor activity of the relevant transporters was evaluated in the in vitro systems used to derive the kinetic parameters applied in our study. Consequently, we utilized transporter expression values reported in literature, which is a common PBK modelling practice (Harwood et al., 2013; Tan et al., 2024). However, this approach increases the model’s uncertainty, because both the expression and activity of transporters can vary between cell-lines, batches, and laboratories (Hayeshi et al., 2008; Harwood et al., 2016; Clerbaux et al., 2018). Future studies should further investigate the active transport of PFAS by *OATs, OATPs* and bile acid-transporters (*BSEP, ASBT, OST*) in in vitro systems. We recommend that the studies also characterize the expression and activity of the transporters of interest, which will enable a more accurate IVIVE of the in vitro-derived parameters.

The NG-PBK model presented in this study predicted observed PFOA concentrations within acceptable accuracy ranges, by implementing the current understanding of the mechanisms driving the toxicokinetics of PFOA. Integration of diseased conditions, or alteration of transporter expression/activity, as previously done by (de Bruijn et al., 2024), (Huang and Isoherranen, 2020), can help in predicting the high inter-individual variability in the toxicokinetics of PFOA that is observed in epidemiological studies. Both the liver and kidneys have been identified as PFOA target organs for PFOA toxicity (Radke et al., 2022; Li et al., 2025; Vujic et al., 2024). Prediction of PFOA concentrations in these organs, can facilitate its NGRA by deriving target-organ-specific health-based guidance values (HBGV). This can be achieved by, integrating quantitative in vitro to in vivo extrapolation (QIVIVE) with PBK modelling. Previously, (Chen et al., 2022; Silva et al., 2024; Iulini et al., 2024) integrated QIVIVE with the EFSA-2020 PBK model to predict the benchmark dose for either liver or immune toxicity based on either plasma, or liver concentrations. However, the uncertainty of the derived HBGV was related to that of the predicted internal concentrations. We believe that the present NG-PBK could be used instead to derive HBGV based on in vitro hepato-/ or nephrotoxicity data, with the application of an uncertainty factor, correcting for the potential under-prediction of liver concentrations derived from the results of our study (Figure S13).

The development of a NG-PBK model poses the foundation for extrapolating the toxicokinetic mechanisms observed for PFOA to other PFAS congeners. A strength of this model lies in the mechanistic determination of tissue-plasma partition coefficients, for which we used experimentally derived distribution-coefficients of PFOA to the different tissue constituents. This approach facilitates the extrapolation of the NG-PBK model to all Perfluoroalkyl Acids (PFAAs) and their alternatives for which these coefficients are already available (Allendorf et al., 2020). Another strength of this model is the introduction of active biliary excretion via BSEP, for which we assumed equal excretion of PFOA as that of the endogenous substrate TCA. This physiologically plausible mechanism of biliary excretion, allows for the extrapolation of this model to other PFAS following the same approach as we did for PFOA.

Following upon these suggestions, and accounting for differences in the binding affinities of PFAS to albumin, globulins, or the transporting proteins (Zhao et al., 2023; Louisse et al., 2024), could help in the extrapolation of the NG-PBK model to other PFAS. Future PFAS-PBK model-development work could also build on the results of (Fischer et al., 2025), who developed a PBK model for different PFAS in mice. The authors observed that inter-congener differences in toxicokinetics were mainly driven by the number of pfcs.For example, they demonstrated that the toxicokinetics of PFAS with more than 7 pfcs were influenced by their permeability across cell membranes, while the toxicokinetics of PFAS with less than 7 pfcs were driven by their active renal excretion.

## 6 Conclusion

To our knowledge, our study is the first to develop a NG-PBK model for PFOA that predicts internal concentrations throughout lifetime. The NG-PBK model mechanistically describes the toxicokinetic processes that contribute to the fate of PFOA in the human body, using only NAM-derived parameters. Most interestingly, our model is the first one that accounts for the changes in albumin concentrations over lifetime and PFOA clearance due to menstruation. We showed that both these mechanisms contribute to the observed sex- and age-differences in the measured serum concentration and half-life of PFOA in humans.

We showed that the NG-PBK model is equivalent to the already available validated PFOA-PBK models and that it more accurately describes the mechanisms driving the toxicokinetics of PFOA in humans. Despite that both the NG-PBK model and previously published models correctly predicted PFOA serum concentrations, our model was not calibrated to observed animal, nor human data. The global sensitivity analysis showed that all the mechanisms described in our model contributed to the total variance of the model outputs. Furthermore, we showed that the extent of the influence of the modelled processes on the predicted PFOA serum concentration and half-life was in agreement with observations in humans, demonstrating the physiological and toxicokinetic correctness of the model.

Our model poses the mechanistic foundation on which future studies can build on, to develop PBK models for other PFAS. Extrapolation of our model to other PFAS, and its integration with HBM data will bring forward the human toxicological risk assessment of PFAS.

## 8 CRediT authorship contribution statement

**Chrysanthi Pachoulide:** Writing – original draft, Writing – review & editing, Methodology, Conceptualization, Software, Investigation, Data curation, Formal analysis, Validation, Visualization. **Carolina Vogs:** Data curation, Visualization, Resources, Writing – review & editing. **Aude Ratier:** Data curation, Visualization, Resources, Writing – review & editing. **Jack Koster:** Software, Writing - review. **Trine Husøy:** Investigation, Writing – review & editing. **Martine Vrijheid:** Resources, Data curation. **Yiyi Xu:** Resources, Data curation. **Antonios Georgelis:** Resources, Data curation. **Karl Forsell:** Resources, Data curation. **Joost Westerhout:** Writing - review & editing, Supervision, Methodology, Software, Project administration, Funding acquisition, Resources. **Nynke I. Kramer:** Writing – review & editing, Supervision, Conceptualisation, Funding acquisition, Resources.

## 10 Acknowledgement

The authors acknowledge the funding from the European Partnership for the Assessment of Risks from Chemicals (PARC project) founded by the European Union’s Horizon Europe research and innovation program under Grant Agreement No 101057014, and the ministries and national agencies of the six countries involved in the funding.

## 11 Appendix A. Supporting information

Supplementary information is available online (LINK)

## 12 Data availability

Model codes and example of input files are available online: (Pachoulide, 2025). The human biomonitoring data that has been used is confidential.

## References

Abraham, K., Mertens, H., Richter, L., Mielke, H., Schwerdtle, T., Monien, B.H., 2024. Kinetics of 15 per- and polyfluoroalkyl substances (pfas) after single oral application as a mixture – a pilot investigation in a male volunteer. Environment International 193, 109047. doi:10.1016/j.envint.2024.109047.

Abraham, K., Mielke, H., Fromme, H., Völkel, W., Menzel, J., Peiser, M., Zepp, F., Willich, S.N., Weikert, C., 2020. Internal exposure to perfluoroalkyl substances (pfass) and biological markers in 101 healthy 1-year-old children: associations between levels of perfluorooctanoic acid (pfoa) and vaccine response. Archives of Toxicology 94, 2131–2147. URL: http://link.springer.com/10.1007/s00204-020-02715-4, doi:10.1007/s00204-020-02715-4.

Al-Majdoub, Z.M., Scotcher, D., Achour, B., Barber, J., Galetin, A., Rostami-Hodjegan, A., 2021. Quantitative proteomic map of enzymes and transporters in the human kidney: Stepping closer to mechanistic kidney models to define local kinetics. Clinical Pharmacology & Therapeutics 110, 1389–1400. doi:10.1002/cpt.2396.

Allendorf, F., Berger, U., Goss, K.U., Ulrich, N., 2019. Partition coefficients of four perfluoroalkyl acid alternatives between bovine serum albumin (bsa) and water in comparison to ten classical perfluoroalkyl acids. Environmental Science: Processes & Impacts 21, 1852–1863. doi:10.1039/c9em00290a.

Allendorf, F., Goss, K.U., Ulrich, N., 2020. Estimating the equilibrium distribution of perfluoroalkyl acids and 4 of their alternatives in mammals. Environmental Toxicology and Chemistry 40, 910–920. doi:10.1002/etc.4954.

Andersson, A.G., Fletcher, T., Xu, Y., Kärrman, A., Pineda, D., Nilsson, C.A., Lindh, C.H., Jakobsson, K., Li, Y., 2025. The relative importance of fecal and urinary excretion of perfluorooctane sulfonic acid and perfluorooctanoic acid after high exposure – an observational study in ronneby, sweden. Environmental Research 285, 122487. doi:10.1016/j.envres.2025.122487.

Bartell, S.M., Calafat, A.M., Lyu, C., Kato, K., Ryan, P.B., Steenland, K., 2010. Rate of decline in serum pfoa concentrations after granular activated carbon filtration at two public water systems in ohio and west virginia. Environmental Health Perspectives 118, 222–228. doi:10.1289/ehp.0901252. citation Key: bartell2010.

Batzella, E., Rosato, I., Pitter, G., Da Re, F., Russo, F., Canova, C., Fletcher, T., 2024. Determinants of pfoa serum half-life after end of exposure: A longitudinal study on highly exposed subjects in the veneto region. Environmental Health Perspectives 132. doi:10.1289/ehp13152.

Biggeri, A., Stoppa, G., Facciolo, L., Fin, G., Mancini, S., Manno, V., Minelli, G., Zamagni, F., Zamboni, M., Catelan, D., Bucchi, L., 2024. All-cause, cardiovascular disease and cancer mortality in the population of a large italian area contaminated by perfluoroalkyl and polyfluoroalkyl substances (1980–2018). Environmental Health 23. doi:10.1186/s12940-024-01074-2.

Bil, W., Zeilmaker, M.J., Bokkers, B.G., 2022. Internal relative potency factors for the risk assessment of mixtures of per- and polyfluoroalkyl substances (pfas) in human biomonitoring. Environmental Health Perspectives 130. doi:10.1289/ehp10009.

Borghese, M.M., Ward, A., MacPherson, S., Manz, K.E., Atlas, E., Fisher, M., Arbuckle, T.E., Braun, J.M., Bouchard, M.F., Ashley-Martin, J., 2024. Serum concentrations of legacy, alternative, and precursor per- and polyfluoroalkyl substances: a descriptive analysis of adult female participants in the mirec-endo study. Environmental Health 23. doi:10.1186/s12940-024-01085-z.

de Bruijn, V.M.P., te Kronnie, W., Rietjens, I.M.C.M., Bouwmeester, H., 2024. Intestinal in vitro transport assay combined with physiologically based kinetic modeling as a tool to predict bile acid levels in vivo. ALTEX - Alternatives to animal experimentation 41, 20–36. doi:10.14573/altex.2302011.

Cao, H., Zhou, Z., Hu, Z., Wei, C., Li, J., Wang, L., Liu, G., Zhang, J., Wang, Y., Wang, T., Liang, Y., 2022. Effect of enterohepatic circulation on the accumulation of per- and polyfluoroalkyl substances: Evidence from experimental and computational studies. Environmental Science & Technology 56, 3214–3224. doi:10.1021/acs.est.1c07176.

Chen, Q., Chou, W.C., Lin, Z., 2022. Integration of toxicogenomics and physiologically based pharmacokinetic modeling in human health risk assessment of perfluorooctane sulfonate. Environmental Science & Technology 56, 3623–3633. doi:10.1021/acs.est.1c06479.

Cheng, W., Ng, C.A., 2017. A permeability-limited physiologically based pharmacokinetic (pbpk) model for perfluorooctanoic acid (pfoa) in male rats. Environmental Science & Technology 51, 9930–9939. doi:10.1021/acs.est.7b02602.

Clerbaux, L.A., Coecke, S., Lumen, A., Kliment, T., Worth, A.P., Paini, A., 2018. Capturing the applicability of in vitro-in silico membrane transporter data in chemical risk assessment and biomedical research. Science of The Total Environment 645, 97–108. doi:10.1016/j.scitotenv.2018.07.122.

Crawford, L., Halperin, S.A., Dzierlenga, M.W., Skidmore, B., Linakis, M.W., Nakagawa, S., Longnecker, M.P., 2023. Systematic review and meta-analysis of epidemiologic data on vaccine response in relation to exposure to five principal perfluoroalkyl substances. Environment International 172, 107734. doi:10.1016/j.envint.2023.107734.

Dahley, C., Goss, K.U., Ebert, A., 2023. Revisiting the pka-flux method for determining intrinsic membrane permeability. European Journal of Pharmaceutical Sciences 191, 106592. doi:10.1016/j.ejps.2023.106592.

Dawson, P.A., Shneider, B.L., Hofmann, A.F., 2006. Bile Formation and the Enterohepatic Circulation. Academic Press, Burlington. chapter 56. pp. 1437–1462. doi:10.1016/B978-012088394-3/50059-3. dOI: 10.1016/B978-012088394-3/50059-3.

Dent, M., Amaral, R.T., Da Silva, P.A., Ansell, J., Boisleve, F., Hatao, M., Hirose, A., Kasai, Y., Kern, P., Kreiling, R., Milstein, S., Montemayor, B., Oliveira, J., Richarz, A., Taalman, R., Vaillancourt, E., Verma, R., Posada, N., Weiss, C., Kojima, H., 2018. Principles underpinning the use of new methodologies in the risk assessment of cosmetic ingredients. Computational Toxicology 7, 20–26. doi:10.1016/j.comtox.2018.06.001.

Ebert, A., Allendorf, F., Berger, U., Goss, K.U., Ulrich, N., 2020. Membrane/water partitioning and permeabilities of perfluoroalkyl acids and four of their alternatives and the effects on toxicokinetic behavior. Environmental Science & Technology 54, 5051–5061. doi:10.1021/acs.est.0c00175.

Ebert, A., Dahley, C., Goss, K.U., 2024. Pitfalls in evaluating permeability experiments with caco-2/mdck cell monolayers. European Journal of Pharmaceutical Sciences 194, 106699. doi:10.1016/j.ejps.2024.106699.

EFSA, 2020. Risk to human health related to the presence of perfluoroalkyl substances in food. EFSA Journal 18. doi:10.2903/j.efsa.2020.6223.

Ehresman, D.J., Froehlich, J.W., Olsen, G.W., Chang, S.C., Butenhoff, J.L., 2007. Comparison of human whole blood, plasma, and serum matrices for the determination of perfluorooctanesulfonate (pfos), perfluorooctanoate (pfoa), and other fluorochemicals. Environmental Research 103, 176–184. doi:10.1016/j.envres.2006.06.008.

Fenton, S.E., Ducatman, A., Boobis, A., DeWitt, J.C., Lau, C., Ng, C., Smith, J.S., Roberts, S.M., 2020. Per- and polyfluoroalkyl substance toxicity and human health review: Current state of knowledge and strategies for informing future research. Environmental Toxicology and Chemistry 40, 606–630. doi:10.1002/etc.4890.

Fischer, F.C., Ludtke, S., Thackray, C., Pickard, H.M., Haque, F., Dassuncao, C., Endo, S., Schaider, L., Sunderland, E.M., 2024. Binding of per- and polyfluoroalkyl substances (pfas) to serum proteins: Implications for toxicokinetics in humans. Environmental Science & Technology 58, 1055–1063. doi:10.1021/acs.est.3c07415.

Fischer, F.C., Thackray, C., Ferguson, N., Chicoine, C., Skende, O., Hu, Z., Zhu, Y., Slitt, A., Sunderland, E.M., 2025. Understanding mechanisms of pfas absorption, distribution, and elimination using a physiologically based toxicokinetic model. Environmental Science & Technology doi:10.1021/acs.est.5c05473.

Fischer, M., Fadda, H.M., 2016. The effect of sex and age on small intestinal transit times in humans. Journal of Pharmaceutical Sciences 105, 682–686. doi:10.1002/jps.24619.

Fu, J., Gao, Y., Cui, L., Wang, T., Liang, Y., Qu, G., Yuan, B., Wang, Y., Zhang, A., Jiang, G., 2016. Occurrence, temporal trends, and half-lives of perfluoroalkyl acids (pfaas) in occupational workers in china. Scientific Reports 6. doi:10.1038/srep38039.

Fujii, Y., Niisoe, T., Harada, K.H., Uemoto, S., Ogura, Y., Takenaka, K., Koizumi, A., 2015. Toxicokinetics of perfluoroalkyl carboxylic acids with different carbon chain lengths in mice and humans. Journal of Occupational Health 57, 1–12. doi:10.1539/joh.14-0136-oa.

Gastellu, T., Karakoltzidis, A., Ratier, A., Bellouard, M., Alvarez, J.C., Le Bizec, B., Rivière, G., Karakitsios, S., Sarigiannis, D.A., Vogs, C., 2025. A comprehensive library of lifetime physiological equations for pbk models: Enhancing dietary exposure modeling with mercury as a case study. Environmental Research 265, 120393. doi:10.1016/j.envres.2024.120393.

Harwood, M.D., Achour, B., Neuhoff, S., Russell, M.R., Carlson, G., Warhurst, G., Rostami-Hodjegan, A., 2016. In vitro–in vivo extrapolation scaling factors for intestinal p-glycoprotein and breast cancer resistance protein: Part ii. the impact of cross-laboratory variations of intestinal transporter relative expression factors on predicted drug disposition. Drug Metabolism and Disposition 44, 476–480. doi:10.1124/dmd.115.067777.

Harwood, M.D., Neuhoff, S., Carlson, G.L., Warhurst, G., Rostami-Hodjegan, A., 2013. Absolute abundance and function of intestinal drug transporters: a prerequisite for fully mechanistic in vitro–in vivo extrapolation of oral drug absorption. Biopharmaceutics & Drug Disposition 34, 2–28. doi:10.1002/bdd.1810.

Hayeshi, R., Hilgendorf, C., Artursson, P., Augustijns, P., Brodin, B., Dehertogh, P., Fisher, K., Fossati, L., Hovenkamp, E., Korjamo, T., Masungi, C., Maubon, N., Mols, R., Müllertz, A., Mönkkönen, J., O’Driscoll, C., Oppers-Tiemissen, H.M., Ragnarsson, E.G.E., Rooseboom, M., Ungell, A.L., 2008. Comparison of drug transporter gene expression and functionality in caco-2 cells from 10 different laboratories. European Journal of Pharmaceutical Sciences 35, 383–396. doi:10.1016/j.ejps.2008.08.004.

Heitmann, P.T., Vollebregt, P.F., Knowles, C.H., Lunniss, P.J., Dinning, P.G., Scott, S.M., 2021. Understanding the physiology of human defaecation and disorders of continence and evacuation. Nature Reviews Gastroenterology & Hepatology 18, 751–769. doi:10.1038/s41575-021-00487-5.

Hong, X., Morgenlander, W.R., Nadeau, K., Wang, G., Frischmeyer-Guerrerio, P.A., Pearson, C., Adams, W.G., Ji, H., Larman, H.B., Wang, X., 2025. Maternal exposure to per- and polyfluoroalkyl substances and epitope level antibody response to vaccines against measles and rubella in children from the boston birth cohort. Environment International 198, 109433. doi:10.1016/j.envint.2025.109433.

Huang, H., Li, X., Deng, Y., San, S., Qiu, D., Xu, A., Luo, J., Xu, L., Li, Y., Zhang, H., Li, Y., 2024. Associations between prenatal exposure to per- and polyfluoroalkyl substances and plasma immune molecules in three-year-old children in china. Toxicology and Applied Pharmacology 490, 117044. doi:10.1016/j.taap.2024.117044.

Huang, W., Isoherranen, N., 2020. Novel mechanistic pbpk model to predict renal clearance in varying stages of ckd by incorporating tubular adaptation and dynamic passive reabsorption. CPT: Pharmacometrics & Systems Pharmacology 9, 571–583. doi:10.1002/psp4.12553.

Husøy, T., Caspersen, I., Thépaut, E., Knutsen, H., Haug, L., Andreassen, M., Gkrillas, A., Lindeman, B., Thomsen, C., Herzke, D., Dirven, H., Wojewodzic, M., 2023. Comparison of aggregated exposure to perfluorooctanoic acid (pfoa) from diet and personal care products with concentrations in blood using a pbpk model – results from the norwegian biomonitoring study in euromix. Environmental Research 239, 117341. doi:10.1016/j.envres.2023.117341.

ICRP, 2002. Basic anatomical and physiological data for use in radiological protection: reference values. Annals of the ICRP 32, 1–277. doi:10.1016/s0146-6453(03)00002-2.

Iulini, M., Russo, G., Crispino, E., Paini, A., Fragki, S., Corsini, E., Pappalardo, F., 2024. Advancing pfas risk assessment: Integrative approaches using agent-based modelling and physiologically-based kinetic for environmental and health safety. Computational and Structural Biotechnology Journal 23, 2763–2778. doi:10.1016/j.csbj.2024.06.036.

Janssen, A.W.F., Duivenvoorde, L.P.M., Beekmann, K., Pinckaers, N., van der Hee, B., Noorlander, A., Leenders, L.L., Louisse, J., van der Zande, M., 2024. Transport of perfluoroalkyl substances across human induced pluripotent stem cell-derived intestinal epithelial cells in comparison with primary human intestinal epithelial cells and caco-2 cells. Archives of Toxicology 98, 3777–3795. doi:10.1007/s00204-024-03851-x.

Johanson, G., Gyllenhammar, I., Ekstrand, C., Pyko, A., Xu, Y., Li, Y., Norström, K., Lilja, K., Lindh, C., Benskin, J.P., Georgelis, A., Forsell, K., Jakobsson, K., Glynn, A., Vogs, C., 2023. Quantitative relationships of perfluoroalkyl acids in drinking water associated with serum concentrations above background in adults living near contamination hotspots in sweden. Environmental Research 219, 115024. doi:10.1016/j.envres.2022.115024.

Karakoltzidis, A., Karakitsios, S.P., Gabriel, C., Sarigiannis, D., 2025. Integrated pbpk modelling for pfoa exposure and risk assessment. Environmental Research 282, 121947. doi:10.1016/j.envres.2025.121947.

Kimura, O., Fujii, Y., Haraguchi, K., Kato, Y., Ohta, C., Koga, N., Endo, T., 2017. Uptake of perfluorooctanoic acid by caco-2 cells: Involvement of organic anion transporting polypeptides. Toxicology Letters 277, 18–23. doi:10.1016/j.toxlet.2017.05.012.

Kinzi, J., Grube, M., Meyer Zu Schwabedissen, H.E., 2021. Oatp2b1 – the underrated member of the organic anion transporting polypeptide family of drug transporters? Biochemical Pharmacology 188, 114534. doi:10.1016/j.bcp.2021.114534.

Li, S., Oliva, P., Zhang, L., Goodrich, J.A., McConnell, R., Conti, D.V., Chatzi, L., Aung, M., 2025. Associations between per-and polyfluoroalkyl substances (pfas) and county-level cancer incidence between 2016 and 2021 and incident cancer burden attributable to pfas in drinking water in the united states. Journal of Exposure Science & Environmental Epidemiology 35, 425–436. doi:10.1038/s41370-024-00742-2.

Li, Y., Fletcher, T., Mucs, D., Scott, K., Lindh, C.H., Tallving, P., Jakobsson, K., 2017. Half-lives of pfos, pfhxs and pfoa after end of exposure to contaminated drinking water. Occupational and Environmental Medicine 75, 46–51. doi:10.1136/oemed-2017-104651.

Li, Z., Xiong, J., 2023. A dynamic inventory database for assessing age-, gender-, and route-specific chronic internal exposure to chemicals in support of human exposome research. Journal of Environmental Management 339, 117867. doi:10.1016/j.jenvman.2023.117867.

Li, Z., Zhang, X., 2023. Assessing human internal exposure to chemicals at different physical activity levels: A physiologically based kinetic (PBK) model incorporating metabolic equivalent of task (MET). Environment International 182, 108312. doi:10.1016/j.envint.2023.108312.

Lin, C.Y., Huey-Jen Hsu, S., Lee, H.L., Wang, C., Sung, F.C., Su, T.C., 2024. Examining a decade-long trend in exposure to per- and polyfluoroalkyl substances and their correlation with lipid profiles: Insights from a prospective cohort study on the young taiwanese population. Chemosphere 364, 143072. doi:10.1016/j.chemosphere.2024.143072.

Lin, J., Chin, S.Y., Tan, S.P.F., Koh, H.C., Cheong, E.J.Y., Chan, E.C.Y., Chan, J.C.Y., 2023. Mechanistic middle-out physiologically based toxicokinetic modeling of transporter-dependent disposition of perfluorooctanoic acid in humans. Environmental Science & Technology 57, 6825–6834. doi:10.1021/acs.est.2c05642.

Lin, P.I.D., Cardenas, A., Hauser, R., Gold, D.R., Kleinman, K.P., Hivert, M.F., Calafat, A.M., Webster, T.F., Horton, E.S., Oken, E., 2021. Per- and polyfluoroalkyl substances and kidney function: Follow-up results from the diabetes prevention program trial. Environment International 148, 106375. doi:10.1016/j.envint.2020.106375.

Loccisano, A.E., Campbell, J.L., Andersen, M.E., Clewell, H.J., 2011. Evaluation and prediction of pharmacokinetics of pfoa and pfos in the monkey and human using a pbpk model. Regulatory Toxicology and Pharmacology 59, 157–175. doi:10.1016/j.yrtph.2010.12.004.

Louisse, J., Dellafiora, L., van den Heuvel, J.J.M.W., Rijkers, D., Leenders, L., Dorne, J.L.C.M., Punt, A., Russel, F.G.M., Koenderink, J.B., 2022. Perfluoroalkyl substances (pfass) are substrates of the renal human organic anion transporter 4 (oat4). Archives of Toxicology 97, 685–696. doi:10.1007/s00204-022-03428-6.

Louisse, J., Pedroni, L., van den Heuvel, J.J., Rijkers, D., Leenders, L., Noorlander, A., Punt, A., Russel, F.G., Koenderink, J.B., Dellafiora, L., 2024. In vitro and in silico characterization of the transport of selected perfluoroalkyl carboxylic acids and perfluoroalkyl sulfonic acids by human organic anion transporter 1 (oat1), oat2 and oat3. Toxicology 509, 153961. doi:10.1016/j.tox.2024.153961.

McNally, K., Cotton, R., Loizou, G.D., 2011. A workflow for global sensitivity analysis of pbpk models. Frontiers in Pharmacology 2. doi:10.3389/fphar.2011.00031.

Mohammadi, F., Rostami, G., Assad, D., Shafiei, M., Hamid, M., Jalaeikhoo, H., 2021. Association of slc22a1, slco1b3 drug transporter polymorphisms and smoking with disease risk and cytogenetic response to imatinib in patients with chronic myeloid leukemia. Laboratory Medicine 52, 584–596. doi:10.1093/labmed/lmab023.

Müller, J., Keiser, M., Drozdzik, M., Oswald, S., 2017. Expression, regulation and function of intestinal drug transporters: an update. Biological Chemistry 398, 175–192. doi:10.1515/hsz-2016-0259.

Nielsen, F., Fischer, F.C., Leth, P.M., Grandjean, P., 2023. Occurrence of major perfluorinated alkylate substances in human blood and target organs. Environmental Science & Technology 58, 143–149. doi:10.1021/acs.est.3c06499.

Niu, S., Cao, Y., Chen, R., Bedi, M., Sanders, A.P., Ducatman, A., Ng, C., 2023. A state-of-the-science review of interactions of per- and polyfluoroalkyl substances (pfas) with renal transporters in health and disease: Implications for population variability in pfas toxicokinetics. Environmental Health Perspectives 131. doi:10.1289/ehp11885.

OECD, 2021a. Guidance document on the characterisation, validation and reporting of physiologically based kinetic (pbk) models for regulatory purposes. OECD Series on Testing and Assessment doi:10.1787/d0de241f-en.

OECD, 2021b. Reconciling terminology of the universe of per- and polyfluoroalkyl substances. OECD Series on Risk Management of Chemicals doi:10.1787/e458e796-en.

Olsen, G.W., Burris, J.M., Ehresman, D.J., Froehlich, J.W., Seacat, A.M., Butenhoff, J.L., Zobel, L.R., 2007. Half-life of serum elimination of perfluorooctanesulfonate,perfluorohexanesulfonate, and perfluorooctanoate in retired fluorochemical production workers. Environmental Health Perspectives 115, 1298–1305. doi:10.1289/ehp.10009.

Oswald, S., 2019. Organic anion transporting polypeptide (oatp) transporter expression, localization and function in the human intestine. Pharmacology & Therapeutics 195, 39–53. doi:10.1016/j.pharmthera.2018.10.007.

Pachoulide, C., 2025. pachrys/PARC_PFOA_NG-PBK: Sprouting (Version 0). Zenodo. doi:10.5281/zenodo.17427866.

Paini, A., Leonard, J.A., Joossens, E., Bessems, J.G.M., Desalegn, A., Dorne, J.L., Gosling, J.P., Heringa, M.B., Klaric, M., Kliment, T., Kramer, N.I., Loizou, G., Louisse, J., Lumen, A., Madden, J.C., Patterson, E.A., Proença, S., Punt, A., Setzer, R.W., Suciu, N., Troutman, J., Yoon, M., Worth, A., Tan, Y.M., 2019. Next generation physiologically based kinetic (ng-pbk) models in support of regulatory decision making. Comput Toxicol 9, 61–72. doi:10.1016/j.comtox.2018.11.002.

Paini, A., Tan, Y.M., Sachana, M., Worth, A., 2021a. Gaining acceptance in next generation pbk modelling approaches for regulatory assessments – an oecd international effort. Computational Toxicology 18, 100163. doi:10.1016/j.comtox.2021.100163.

Paini, A., Worth, A., Kulkarni, S., Ebbrell, D., Madden, J., 2021b. Assessment of the predictive capacity of a physiologically based kinetic model using a read-across approach. Computational Toxicology 18, 100159. doi:10.1016/j.comtox.2021.100159.

Pletz, J., Allen, T.J., Madden, J.C., Cronin, M.T., Webb, S.D., 2021. A mechanistic model to study the kinetics and toxicity of salicylic acid in the kidney of four virtual individuals. Computational Toxicology 19, 100172. doi:10.1016/j.comtox.2021.100172.

Poothong, S., Papadopoulou, E., Padilla-Sánchez, J.A., Thomsen, C., Haug, L., 2020. Multiple pathways of human exposure to poly- and perfluoroalkyl substances (pfass): From external exposure to human blood. Environment International 134, 105244. doi:10.1016/j.envint.2019.105244.

Pottel, H., Hoste, L., Dubourg, L., Ebert, N., Schaeffner, E., Eriksen, B., Melsom, T., Lamb, E.J., Rule, A.D., Turner, S.T., Glassock, R.J., De Souza, V., Selistre, L., Mariat, C., Martens, F., Delanaye, P., 2016. An estimated glomerular filtration rate equation for the full age spectrum. Nephrology Dialysis Transplantation 31, 798–806. doi:10.1093/ndt/gfv454.

Poulin, P., Haddad, S., 2018. Extrapolation of the hepatic clearance of drugs in the absence of albumin in vitro to that in the presence of albumin in vivo : Comparative assessement of 2 extrapolation models based on the albumin-mediated hepatic uptake theory and limitations and mechanistic insights. Journal of Pharmaceutical Sciences 107, 1791–1797. doi:10.1016/j.xphs.2018.03.012.

Punt, A., Baltazar, M.T., Nicol, B., Cable, S., Hewitt, N.J., Cubberley, R., Spriggs, S., Dent, M.P., Li, H., 2025. Building confidence in pbk model predictions in the absence of human kinetic data: Benzophenone-4 case study. ALTEX - Alternatives to Animal Experimentation URL: https://www.altex.org/index.php/altex/article/view/2933, doi:10.14573/altex.2501211.

Radke, E.G., Wright, J.M., Christensen, K., Lin, C.J., Goldstone, A.E., Lemeris, C., Thayer, K.A., 2022. Epidemiology evidence for health effects of 150 per- and polyfluoroalkyl substances: A systematic evidence map. Environmental Health Perspectives 130. doi:10.1289/ehp11185.

Ragnarsdóttir, O., Abou-Elwafa Abdallah, M., Harrad, S., 2024. Dermal bioavailability of perfluoroalkyl substances using in vitro 3d human skin equivalent models. Environment International 188, 108772. doi:10.1016/j.envint.2024.108772.

Ratier, A., Casas, M., Grazuleviciene, R., Slama, R., Småstuen Haug, L., Thomsen, C., Vafeiadi, M., Wright, J., Zeman, F.A., Vrijheid, M., Brochot, C., 2024. Estimating the dynamic early life exposure to pfoa and pfos of the helix children: Emerging profiles via prenatal exposure, breastfeeding, and diet. Environment International 186, 108621. doi:10.1016/j.envint.2024.108621.

Richterová, D., Govarts, E., Fábelová, L., Rausová, K., Rodriguez Martin, L., Gilles, L., Remy, S., Colles, A., Rambaud, L., Riou, M., Gabriel, C., Sarigiannis, D., Pedraza-Diaz, S., Ramos, J., Kosjek, T., Snoj Tratnik, J., Lignell, S., Gyllenhammar, I., Thomsen, C., Haug, L., Kolossa-Gehring, M., Vogel, N., Franken, C., Vanlarebeke, N., Bruckers, L., Stewart, L., Sepai, O., Schoeters, G., Uhl, M., Castaño, A., Esteban López, M., Göen, T., Palkovičová Murínová, Ľ., 2023. Pfas levels and determinants of variability in exposure in european teenagers – results from the hbm4eu aligned studies (2014–2021). International Journal of Hygiene and Environmental Health 247, 114057. doi:10.1016/j.ijheh.2022.114057.

Rosato, I., Bonato, T., Fletcher, T., Batzella, E., Canova, C., 2024. Estimation of per- and polyfluoroalkyl substances (pfas) half-lives in human studies: a systematic review and meta-analysis. Environmental Research 242, 117743. doi:10.1016/j.envres.2023.117743.

Rosato, I., Zare Jeddi, M., Ledda, C., Gallo, E., Fletcher, T., Pitter, G., Batzella, E., Canova, C., 2022. How to investigate human health effects related to exposure to mixtures of per- and polyfluoroalkyl substances: A systematic review of statistical methods. Environmental Research 205, 112565. doi:10.1016/j.envres.2021.112565.

Ruark, C.D., Song, G., Yoon, M., Verner, M., Andersen, M.E., Clewell, H.J., Longnecker, M.P., 2017. Quantitative bias analysis for epidemiological associations of perfluoroalkyl substance serum concentrations and early onset of menopause. Environment International 99, 245–254. doi:10.1016/j.envint.2016.11.030.

Ryu, S., Yamaguchi, E., Sadegh Modaresi, S.M., Agudelo, J., Costales, C., West, M.A., Fischer, F., Slitt, A.L., 2024. Evaluation of 14 pfas for permeability and organic anion transporter interactions: Implications for renal clearance in humans. Chemosphere 361, 142390. doi:10.1016/j.chemosphere.2024.142390.

Saltelli, A., Tarantola, S., Chan, K.P.S., 1999. A quantitative model-independent method for global sensitivity analysis of model output. Technometrics 41, 39–56. doi:10.1080/00401706.1999.10485594.

Sayyed, K., Le Vee, M., Abdel-Razzak, Z., Fardel, O., 2017. Inhibition of organic anion transporter (oat) activity by cigarette smoke condensate. Toxicology in Vitro 44, 27–35. doi:10.1016/j.tiv.2017.06.014.

Sayyed, K., Vee, M.L., Abdel-Razzak, Z., Jouan, E., Stieger, B., Denizot, C., Parmentier, Y., Fardel, O., 2016. Alteration of human hepatic drug transporter activity and expression by cigarette smoke condensate. Toxicology 363-364, 58–71. doi:10.1016/j.tox.2016.07.011.

Schultz, A.A., Stanton, N., Shelton, B., Pomazal, R., Lange, M.A., Irving, R., Meiman, J., Malecki, K.C., 2023. Biomonitoring of perfluoroalkyl and polyfluoroalkyl substances (pfas) from the survey of the health of wisconsin (show) 2014–2016 and comparison with the national health and nutrition examination survey (nhanes). Journal of Exposure Science & Environmental Epidemiology 33, 766–777. doi:10.1038/s41370-023-00593-3.

Scotcher, D., Jones, C., Rostami-Hodjegan, A., Galetin, A., 2016. Novel minimal physiologically-based model for the prediction of passive tubular reabsorption and renal excretion clearance. European Journal of Pharmaceutical Sciences 94, 59–71. doi:10.1016/j.ejps.2016.03.018.

Seals, R., Bartell, S.M., Steenland, K., 2011. Accumulation and clearance of perfluorooctanoic acid (pfoa) in current and former residents of an exposed community. Environmental Health Perspectives 119, 119–124. doi:10.1289/ehp.1002346.

Shirke, A.V., Radke, E.G., Lin, C., Blain, R., Vetter, N., Lemeris, C., Hartman, P., Hubbard, H., Angrish, M., Arzuaga, X., Congleton, J., Davis, A., Dishaw, L.V., Jones, R., Judson, R., Kaiser, J.P., Kraft, A., Lizarraga, L., Noyes, P.D., Patlewicz, G., Taylor, M., Williams, A.J., Thayer, K.A., Carlson, L.M., 2024. Expanded systematic evidence map for hundreds of per- and polyfluoroalkyl substances (pfas) and comprehensive pfas human health dashboard. Environmental Health Perspectives 132. doi:10.1289/ehp13423.

Silva, A., Loizou, G.D., McNally, K., Osborne, O., Potter, C., Gott, D., Colbourne, J.K., Viant, M.R., 2024. A novel method to derive a human safety limit for pfoa by gene expression profiling and modelling. Frontiers in Toxicology 6, 1368320. doi:10.3389/ftox.2024.1368320.

Smeltz, M., Wambaugh, J.F., Wetmore, B.A., 2023. Plasma protein binding evaluations of per- and polyfluoroalkyl substances for category-based toxicokinetic assessment. Chemical Research in Toxicology 36, 870–881. doi:10.1021/acs.chemrestox.3c00003.

Sugano, K., Kansy, M., Artursson, P., Avdeef, A., Bendels, S., Di, L., Ecker, G.F., Faller, B., Fischer, H., Gerebtzoff, G., Lennernaes, H., Senner, F., 2010. Coexistence of passive and carrier-mediated processes in drug transport. Nature Reviews Drug Discovery 9, 597–614. doi:10.1038/nrd3187.

Sunderland, E.M., Hu, X.C., Dassuncao, C., Tokranov, A.K., Wagner, C.C., Allen, J.G., 2018. A review of the pathways of human exposure to poly- and perfluoroalkyl substances (pfass) and present understanding of health effects. Journal of Exposure Science & Environmental Epidemiology 29, 131–147. doi:10.1038/s41370-018-0094-1.

Tan, S.P.F., Tillmann, A., Murby, S.J., Rostami-Hodjegan, A., Scotcher, D., Galetin, A., 2024. Albumin-mediated drug uptake by organic anion transporter 1/3 is real: Implications for the prediction of active renal secretion clearance. Molecular Pharmaceutics 21, 4603–4617. doi:10.1021/acs.molpharmaceut.4c00504.

Thompson, C.V., Webb, S.D., Leedale, J.A., Penson, P.E., Paini, A., Ebbrell, D., Madden, J.C., 2024. Using read-across to build physiologically-based kinetic models: Part 2. case studies for atenolol and flumioxazin. Computational Toxicology 29, 100293. doi:10.1016/j.comtox.2023.100293.

Tiburtini, G.A., Bertarini, L., Bersani, M., Dragani, T.A., Rolando, B., Binello, A., Barge, A., Spyrakis, F., 2024. In silico prediction of the interaction of legacy and novel per- and poly-fluoroalkyl substances (pfas) with selected human transporters and of their possible accumulation in the human body. Archives of Toxicology 98, 3035–3047. doi:10.1007/s00204-024-03797-0.

Tojo, A., Kinugasa, S., 2012. Mechanisms of glomerular albumin filtration and tubular reabsorption. International Journal of Nephrology 2012, 1–9. doi:10.1155/2012/481520.

Upson, K., Shearston, J.A., Kioumourtzoglou, M.A., 2022. An epidemiologic review of menstrual blood loss as an excretion route for per- and polyfluoroalkyl substances. Current Environmental Health Reports 9, 29–37. doi:10.1007/s40572-022-00332-0.

Utsey, K., Gastonguay, M.S., Russell, S., Freling, R., Riggs, M.M., Elmokadem, A., 2020. Quantification of the impact of partition coefficient prediction methods on physiologically based pharmacokinetic model output using a standardized tissue composition. Drug Metabolism and Disposition 48, 903– 916. doi:10.1124/dmd.120.090498.

Vujic, E., Ferguson, S.S., Brouwer, K.L.R., 2024. Effects of pfas on human liver transporters: implications for health outcomes. Toxicological Sciences 200, 213–227. doi:10.1093/toxsci/kfae061.

Weaving, G., Batstone, G.F., Jones, R.G., 2015. Age and sex variation in serum albumin concentration: an observational study. Annals of Clinical Biochemistry: International Journal of Laboratory Medicine 53, 106–111. doi:10.1177/0004563215593561.

WHO, 2010. Characterization and application of physiologically based pharmacokinetic models in risk assessment. Technical Report. International Program on Chemical Safety (IPCS), Inter-Organization Program for the Sound Management of Chemicals (IOMC), World Health Organisation (WHO). URL: https://www.who.int/publications/i/item/9789241500906.

Willmann, S., Schmitt, W., Keldenich, J., Lippert, J., Dressman, J.B., 2004. A physiological model for the estimation of the fraction dose absorbed in humans. Journal of Medicinal Chemistry 47, 4022–4031. doi:10.1021/jm030999b.

Worley, R.R., Moore, S.M., Tierney, B.C., Ye, X., Calafat, A.M., Campbell, S., Woudneh, M.B., Fisher, J., 2017. Per- and polyfluoroalkyl substances in human serum and urine samples from a residentially exposed community. Environment International 106, 135–143. doi:10.1016/j.envint.2017.06.007.

Wu, H., Yoon, M., Verner, M., Xue, J., Luo, M., Andersen, M.E., Longnecker, M.P., Clewell, H.J., 2015. Can the observed association between serum perfluoroalkyl substances and delayed menarche be explained on the basis of puberty-related changes in physiology and pharmacokinetics? Environment International 82, 61–68. doi:10.1016/j.envint.2015.05.006.

Xu, Y., Fletcher, T., Pineda, D., Lindh, C.H., Nilsson, C., Glynn, A., Vogs, C., Norström, K., Lilja, K., Jakobsson, K., Li, Y., 2020. Serum half-lives for short- and long-chain perfluoroalkyl acids after ceasing exposure from drinking water contaminated by firefighting foam. Environmental Health Perspectives 128. doi:10.1289/ehp6785.

Yang, C.H., Glover, K.P., Han, X., 2010. Characterization of cellular uptake of perfluorooctanoate via organic anion-transporting polypeptide 1a2, organic anion transporter 4, and urate transporter 1 for their potential roles in mediating human renal reabsorption of perfluorocarboxylates. Toxicological Sciences 117, 294–302. doi:10.1093/toxsci/kfq219.

Zhang, Y., Beesoon, S., Zhu, L., Martin, J.W., 2013. Biomonitoring of perfluoroalkyl acids in human urine and estimates of biological half-life. Environmental Science & Technology 47, 10619–10627. doi:10.1021/es401905e.

Zhao, L., Teng, M., Zhao, X., Li, Y., Sun, J., Zhao, W., Ruan, Y., Leung, K.M., Wu, F., 2023. Insight into the binding model of per- and polyfluoroalkyl substances to proteins and membranes. Environment International 175, 107951. doi:10.1016/j.envint.2023.107951.

