## Supplemental Material for "A Human Next Generation PBK Model for PFOA"

### Supplementary Information
#### A Human Next Generation PBK Model for PFOA

Chrysanthi Pachoulide[1, *], Carolina Vogs[2], Aude Ratier[3,4], Jack Koster[5], Trine Husøy[6], Martine Vrijheid[7], Yiyi Xu[8], Antonios Georgelis[9], Karl Forsell[10], Joost Westerhout[5], and Nynke I. Kramer[1]

[1]Wageningen University and Research, Division of Toxicology, Wageningen
[2]Swedish University of Agricultural Sciences, Department of Animal Biosciences, Uppsala
[3]French National Institute for Industrial Environment and Risks (INERIS), Unit of Experimental Toxicology and Modelling, Verneuil-en-Halatte
[4]Université de Picardie Jules Verne, PériTox Laboratory, UMR-I 01 INERIS, Amiens
[5]National Institute for Public Health and the Environment (RIVM), Center for Prevention, Lifestyle and Health, Bilthoven
[6]Norwegian institute of Public Health, Department of Food Safety, Oslo
[7]Universitat Pompeu Fabra, ISGlobal, Barcelona
[8]University of Gothenburg, School of Public Health and Community Medicine, Gothenburg
[9]Karolinska Institutet, Institute of Environmental Medicine, Stockholm
[10]Region Vasterbotten, Umeå

### 1 PBK model parameters

#### 1.1 Physiological parameters

##### 1.1.1 Serum albumin over lifetime

Serum albumin concentration was dynamically calculated over lifetime. For this, a polynomial equation was derived, using the reported average serum albumin values in males and females throughout lifetime reported by (Weaving et al.,

2015). This led to the below equations for males Equation S1 and females Equation S2 :

$$S_{alb} = 3.9330e^{+01} + (5.8156e^{-01} \times age) - (1.9072e^{-02} \times age^2) + (2.3220e^{-0.4} \times age^3) - (1.0313e^{-06} \times age^4) \tag{S1}$$

$$S_{alb} = 4.0540e^{+01} + (6.0605^{-01} \times age) - (3.5316^{-02} \times age^2) + (7.9681^{-0.4} \times age^3) - (7.8062^{-06} \times age^4) + (2.7227e^{-08} \times age^5) \tag{S2}$$

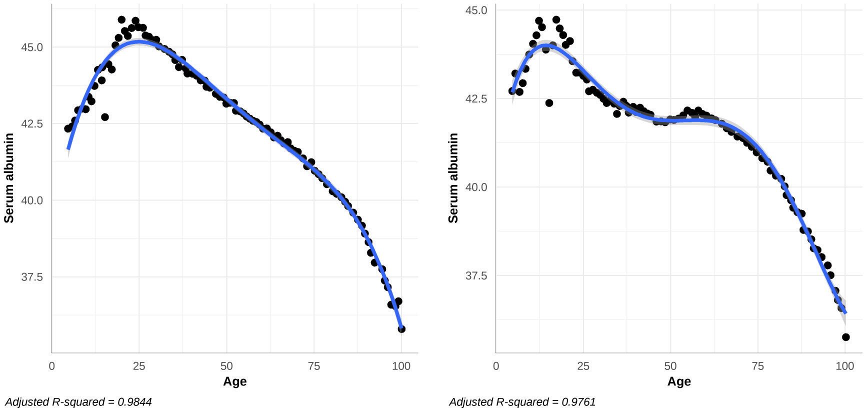


Figure S1: Modelled serum albumin concentration over lifetime in males (left) and females (right).

###### 1.1.2 Glomerular filtration rate over lifetime

Traditionally, the GFR is calculated based on the filtration fraction of 0.18 from the kidney plasma flow (Valentin, 2002). Given the sensitivity of the model to this parameter, we updated the standard equation for estimating GFR with the FAS equation (Pottel et al., 2016), developed based on studies including 6870 individuals (735 children, and 1764 adults above 70 years of age), as also described in Materials and Methods.

##### 1.2 PFOA tissue-plasma partition coefficients

Previous models (Panel), 2020; Loccisano et al., 2011) used tissue-plasma partition coefficients (Kps) taken from a rat toxicokinetic study (Kudo et al., 2007). However, we consider this approach uncertain, due to the physiological differences

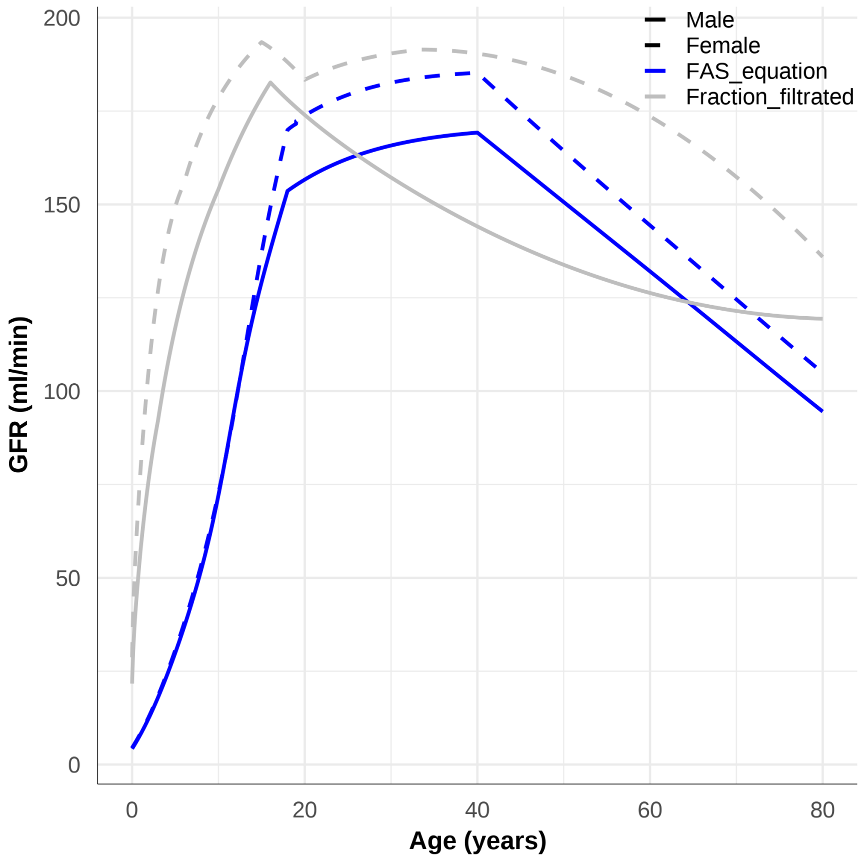


Figure S2: GFR depending on the estimation method. Blue and grey lines represent the estimated GFR over lifetime, using the FAS equation and a defined fraction fitrated of the kidney blood flow. Females and males are represented in solid and dashed lines.

between rats and humans. Other authors, (Ratier et al., 2024; Karakoltzidis et al., 2025), where available, used Kps derived from healthy human forensic tissue autopsies (Maestri et al., 2006; Nielsen et al., 2023) for their PFOA PBK models. This approach is more physiologically relevant; nevertheless, as discussed by (Nielsen et al., 2023), the tissue sample sizes were limited and may not have been representative of the PFOA concentration in the entire tissue. Furthermore, (Nielsen et al., 2023) observed variability in blood composition due to blood coagulation in some of the samples studied. Consequently, and due to the potential disparities in post-mortem tissue perfusion (leading to higher perfusion of certain tissues compared to others), we decided not to use these partition coefficients.

Furthermore, the utilization of experimentally derived partition coefficients was not undertaken due to their specificity to PFOA, and their lack of mechanistic underpinnings. The application of these partition coefficients would have hindered the extrapolation of the PBK model to other PFAS, given that tissue distribution is compound-specific and contingent on the physicochemical properties of each compound. Observations of discrepancies in the tissue distribution of perfluoroalkyl substances (PFAS) were indeed made by (Nielsen et al., 2023). The authors observed that the distribution of PFOA and PFHxS seemed to be driven by active transport, while that of PFNA and PFOS was driven by tissue binding.

Alternatively, Kps, are definable by mechanistic tissue-plasma partition algorithms (Rodgers et al., 2005; Rodgers and Rowland, 2006, 2007; Poulin et al., 2001; Peyret et al., 2010; Utsey et al., 2020). This approach uses physiochemical properties to predict the affinities of a chemical to proteins or lipids (Figure S3). It assumes that these depend on the lipophilicity and ionisation status of the chemical. Consequently, the final steady-state tissue-plasma distribution is driven by the chemical's sorption capacity to proteins and lipids and the differences in protein and lipid content in tissues (Rodgers and Rowland, 2007; Utsey et al., 2020; Peyret et al., 2010). This mechanistic approach is especially useful in the development of NG-PBK models as it relies only on physicochemical properties which are readily available for most chemicals.

Nonetheless, this approach could not be applied for PFAS, because they fall outside the chemical applicability domain of these algorithms. Moreover, there is considerable uncertainty in the physicochemical properties of PFAS, that are needed to calculate Kps. For example, there is high variability in experimentally and in silico derived acid dissociation constant (pKa) for PFOA, with values ranging from 0.2 to 3.8. Furthermore, the octanol-water partition coefficient (Kow) may not be a relevant descriptor, because PFOA sits at the interface of water and octanol, given that it is both hydrophilic and hydrophobic.

In this study, Kps were calculated using the sorption coefficients of PFOA to storage lipids, membrane lipids, albumin, structural proteins and fatty acid binding protein (FABP) derived by (Allendorf et al., 2021), and as shown in Figure S4, following equation 3. Physiological data about the relevant tissue components were taken from (Allendorf et al., 2021) and (Utsey et al., 2020) and are found in Table S1. The predicted partition coefficients were compared with those used in previous PBK models for PFOA. To enable this comparison, we predicted the Kps for both human and rats.

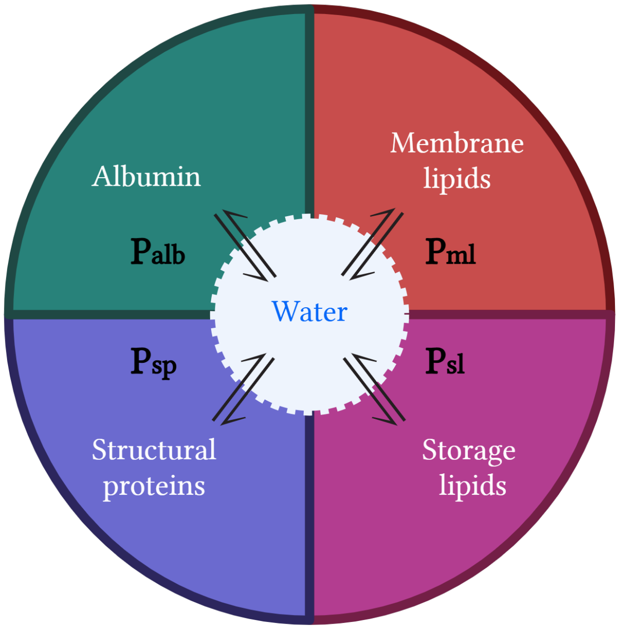


Figure S3: Schematic representation of the partitioning of PFOA between the different tissue constituents. Equilibrium arrows illustrate that the amount of PFOA is in equilibrium between the different tissue constituents depending on its partition coefficients (P) in albumin (Palb), membrane lipids (Pml), structural proteins (Psp) and storage lipids (Psl).

As shown in Figure S5 the partition coefficients calculated in this paper are very similar to those calculated in the reference paper (Allendorf et al., 2021). As illustrated in Figure S6 significant differences between the predicted human and rat Kps were only observed for the adipose and skin tissues, while all other tissues had similar Kps for both sexes. However, as demonstrated in Figure S7, there are discrepancies between the Kps derived in this study and those used in previous PFOA-PBK models (Karakoltzidis et al., 2025; Panel), 2020; Loccisano et al., 2011; Ratier et al., 2024). Importantly, it can be observed the Kps predicted in this study were systematically smaller than those used in previous models and this discrepancy was most remarkable for the liver, kidney and gut. It can also be observed that the liver-plasma Kp used by EFSA and derived from (Kudo et al., 2007) was much higher compared to that used by this study and Karakoltzidis et al. (2025). Further evaluation of these coefficients against in vivo human and rat data was performed in the original paper (Allendorf et al.,

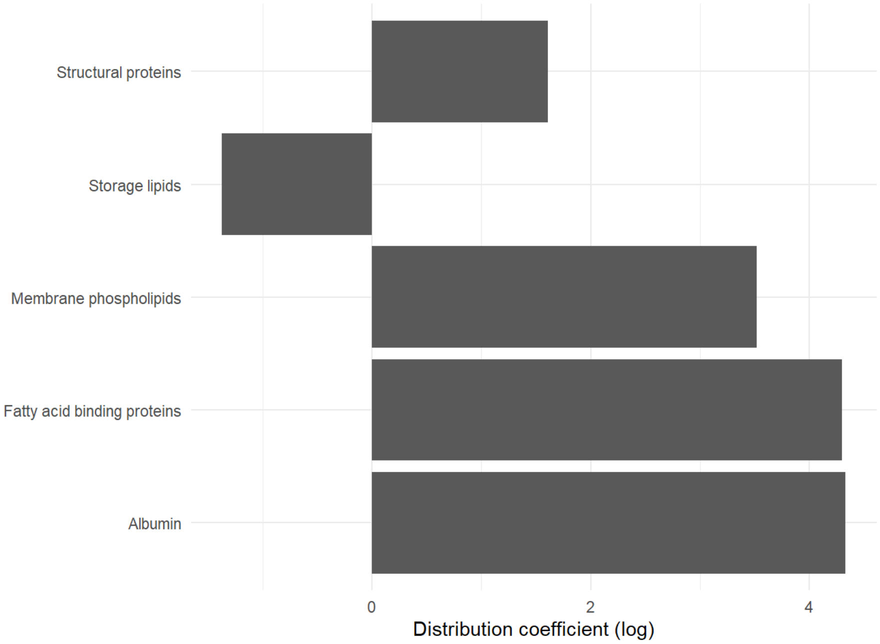


Figure S4: PFOA distribution coefficients to the different tissue constituents, as determined experimentally by (Allendorf et al., 2019; Ebert et al., 2020; Allendorf et al., 2021). Distribution coefficients (tissue constituent/water) are shown on a log scale. Tissue constituents were: structural proteins (actin and myosin extracted from chicken breast fillet), storage lipids (unsaturated fatty acids found in olive oil), membrane lipids (1-palitoyl-2oleoyl-glycero-3-phosphochiline (POPC) found in liposomes, fatty acid binding proteins, and albumin (bovine serum albumin, fatty acid free).

Table S1: Tissue Composition in humans; "f-SL" fractional volume of storage lipids, equals to the volume of total fat minus total phospholipids (neutral + acidic); "f-LM" fractional volume of membrane lipids, equals to the volume of the neutral phospholipids; "f-ALB" fractional volume of albumin, equals to the volume of albumin in the interstitial space; "f-SP" fractional volume of structural proteins, equals to the volume of actine and myosin; "f-W" fractional volume of water; "f-FABP" fractional volume of fatty acid binding protein.

| Tissue | f-SL | f_ML | f_ALB | f_SP | f_W | f_FABP |
|---|---|---|---|---|---|---|
| Adipose | 0.798 | 0.0478 | 0.0009 | 0.0500 | 0.153 | 0.0000 |
| Brain | 0.045 | 0.0553 | 0.0000 | 0.0800 | 0.771 | 0.0000 |
| Stomach | 0.047 | 0.0126 | 0.0007 | 0.1300 | 0.790 | 0.0000 |
| Intestine | 0.047 | 0.0126 | 0.0007 | 0.1300 | 0.790 | 0.0000 |
| Heart | 0.089 | 0.0079 | 0.0009 | 0.1700 | 0.742 | 0.0000 |
| Kidney | 0.036 | 0.0166 | 0.0014 | 0.1667 | 0.778 | 0.0033 |
| Liver | 0.037 | 0.0115 | 0.0012 | 0.1769 | 0.737 | 0.0031 |
| Lung | 0.003 | 0.0056 | 0.0027 | 0.1800 | 0.796 | 0.0000 |
| Muscle | 0.013 | 0.0092 | 0.0009 | 0.1700 | 0.773 | 0.0000 |
| Skin | 0.036 | 0.0502 | 0.0018 | 0.2900 | 0.667 | 0.0000 |
| Spleen | 0.014 | 0.0103 | 0.0013 | 0.1900 | 0.783 | 0.0000 |
| Blood | 0.003 | 0.0039 | 0.0176 | 0.1624 | 0.807 | 0.0000 |
| Plasma | 0.003 | 0.0050 | 0.0294 | 0.0221 | 0.928 | 0.0000 |
| Gonads | 0.000 | 0.0107 | 0.0008 | 0.1200 | 0.810 | 0.0000 |
| Bone | 0.074 | 0.0016 | 0.0000 | 0.2680 | 0.446 | 0.0000 |

2021) and was therefore not repeated here.

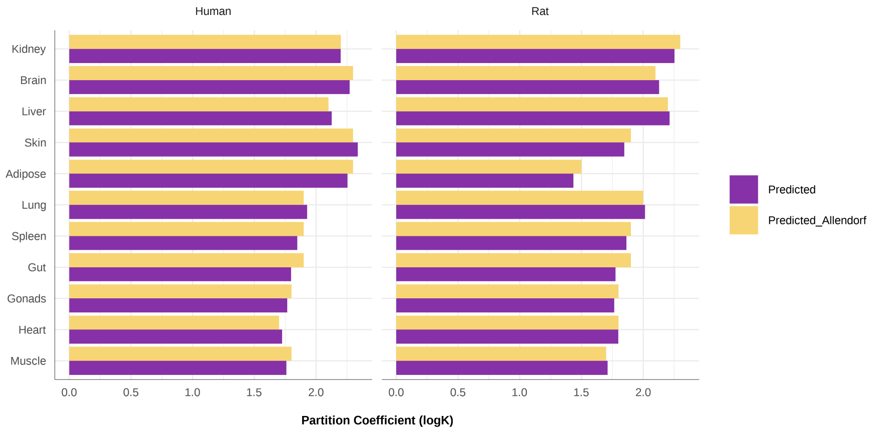


Figure S5: Comparison of the predicted tissue:plasma distribution coefficients in this study, with those predicted by (Allendorf et al., 2021). Distribution coefficients are presented in logarithmic scale. Contrary to partition coefficients, distribution coefficients are not corrected for the fraction unbound in plasma.

#### 1.3 Active transport

PFOA has been shown to be a substrate of transporters from both the organic anion transporter (OAT) and organic anion transporting polypeptide (OATP) families, with different affinities and transport velocities depending on the transporter isoform (Kimura et al., 2017; Lin et al., 2023; Ruggiero et al., 2021; Louisse et al., 2022, 2024). OAT and OATP transporters are expressed in the epithelium that separate blood from tissues and are ubiquitous in the human body (Nigam and Granados, 2023). We focus here on the intestine, kidney and liver, which are the excreting organs of PFOA. The active transport of PFOA in intestine, kidney and liver was studied in in vitro experiments using the Caco-2 (colorectal adenocarcinoma), CHO (Chinese hamster ovary), HEK293 (human embryonic kidney) and MDCKII-LE (Madin-Darby canine kidney) cell lines. While the Caco-2 and MDCKII-LE cell lines were used specifically to study its transport in the intestine and kidney respectively (Kimura et al., 2017; Janssen et al., 2024; Ryu et al., 2024), the other cell lines were used to model both the kidney and liver (Ruggiero et al., 2021; Yang et al., 2010, Lin et al. (2023); Louisse et al., 2022; Louisse et al., 2024). This was possible because in each study the respective transporter was over-expressed in the cell system.

To accurately model active transport processes, PBK modelers have developed permeability-limited PBK models. In these models, each organ is divided into extra cellular (including the vascular and interstitial space) and intra cellular compartments. Permeability-limited PBK models hypothesize that the organ

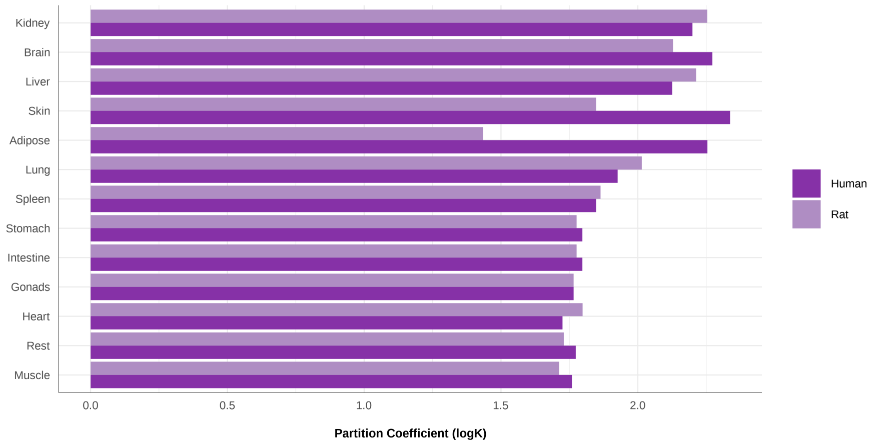


Figure S6: Comparison of tissue:plasma partition distribution coefficients in human and rat predicted by this study. Distribution coefficients are presented in logarithmic scale. Contrary to partition coefficients, distribution coefficients are not corrected for the fraction unbound in plasma.

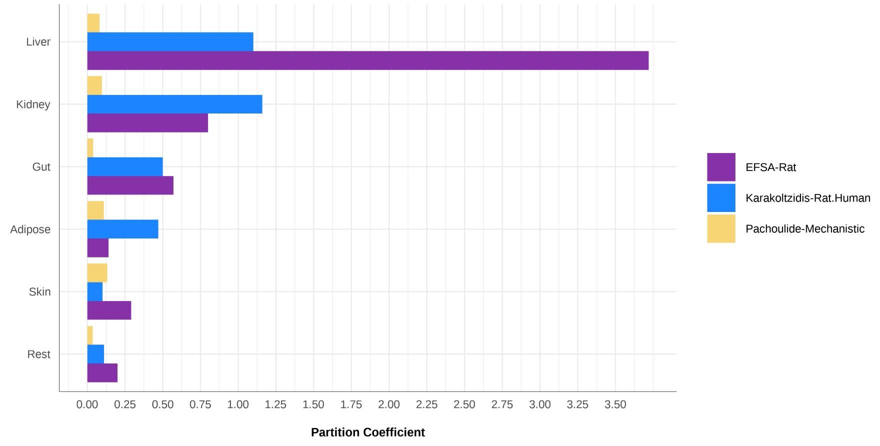


Figure S7: Comparison of the tissue-plasma partition coefficients derived in this study from those used in previously published PFOA PBK models. Partition coefficients were corrected for the fraction unbound in plasma.

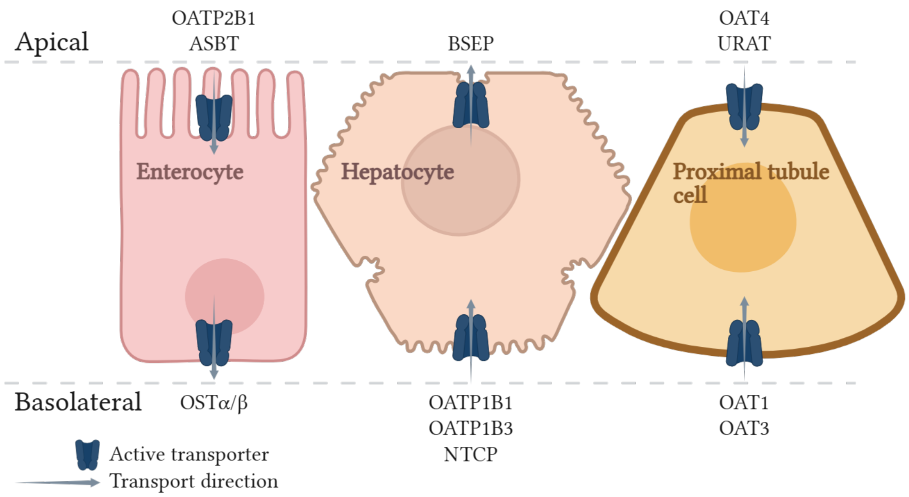


Figure S8: Summary of the active transporters that are potentially involved in the influx, or efflux of PFOA to and from the enterocytes, hepatocytes and proximal tubule cells (Louisse et al., 2022, 2024; Ryu et al., 2024; Vujic et al., 2024; Cao et al., 2022; Niu et al., 2023; Tiburtini et al., 2024; Zhao et al., 2023; Lin et al., 2023; Kimura et al., 2017).

distribution of a chemical is limited by its ability to permeate cell membranes. Such permeability-limited models have recently been developed by (Lin et al., 2023) and (Karakoltzidis et al., 2025) for PFOA in humans and by (Fischer et al., 2025) in rats.

All studies described a permeability-limited distribution of PFOA to the liver and kidney. The authors divided the tissues into vascular, extra vascular and cellular compartments, and assumed that PFOA can move between these compartments via passive diffusion and active transport. However, both human models had to introduce a fitted efflux transport clearance from both the liver and kidney cellular compartments to the vascular compartments in order to accurately describe PFOA serum concentrations (Lin et al., 2023; Karakoltzidis et al., 2025). This suggests that limiting the distribution of PFOA between the vascular and extracellular compartment to an active process underestimates the movement, and seemingly rapid equilibrium of PFOA between these spaces.

In this study we applied a semi-permeability approach to model the kidney and intestine, which describes the current knowledge regarding PFOA distribution to these tissues. We employed the term 'semi-permeability' to denote the assumption of an instant equilibrium between the vascular, interstitial and cellular compartments of the tissue, with active transport contributing to the transfer of PFOA from the organ tissue to the intracellular compartment. Therefore, in this case, distribution of PFOA to the organs from plasma was assumed to be perfusion-limited. An argument supporting the assumptions of a perfusion-limited distribution from plasma, and of an instant equilibrium between the vascular, extracellular and intracellular compartments of the intestine and kidney, is the high membrane permeability of PFOA (Ebert et al., 2020). Passive diffusion is determined by the effective permeability and the environment pH at each side of the membrane, as only the available neutral species can permeate across the lipid bilayer of a membrane. The fraction of neutral species available in the corresponding environment depends on the pH. PFOA has a low pKa and therefore, the more acidic the environment, the more neutral species are available to permeate. This was demonstrated in membrane/water partitioning studies where the effective permeability of PFOA was higher in the direction of intracellular to vascular space because the pH is lower intracellularly than vascularly (Ebert et al., 2020). At pH 7 (pH of intracellular space), PFOA had an effective permeability coefficient (Peff) of 1.3e-4 cm/s, while at pH 7.4 (pH of plasma and extracellular water), it had a Peff of 5.0e-5 cm/s (Ebert et al., 2020).

#### 1.4 Oral and dermal uptake

Intestinal and dermal uptake were modelled as uptake clearances from the intestinal lumen to the intestinal tissue and the skin barrier to the skin tissue, using Papp values obtained *in vitro* by (Janssen et al., 2024) and (Ragnarsdóttir et al., 2024) respectively. We scaled $P_{app}$ values to in vivo uptake clearances, in L/day, following Equation S3, where $SA_{matix}$ stands for surface area of the concerned matrix (small intestinal lumen, or skin barrier).

$$CL_{uptake} = P_{app} * SA_{matrix} \tag{S3}$$

#### 1.5 Enterohepatic circulation

The mechanisms driving the enterohepatic circulation (EHC) and faecal elimination of PFOA are: uptake from the intestine, active uptake from the portal blood to the hepatocytes and active excretion from the hepatocytes to the bile. To best describe these mechanisms, we developed a semi-permeability-limited intestine, which was split into lumen and tissue (including plasma), and a permeability limited-liver, which was split into the intracellular and extracellular (plasma and interstitial fluids).

Model equations, describe the following steps of EHC and faecal elimination:

1. In the intestinal lumen PFOA can either be eliminated to the faeces following the physiological intestinal bowel transit time, or be taken up to the intestinal tissue via both passive diffusion and active transport by OATP2B1 Equation S4

$$\frac{dA_{IL}}{dt} = OD - t_{co}A_{IL} - CL_{IL}C_{IL} + A_{transported_{BSEP}} - A_{transported_{OATP2B1}} \tag{S4}$$

2. From the intestinal tissue PFOA follows the portal plasma flow to the extracellular space of the liver Equation S5

$$\frac{dA_I}{dt} = Q_I(CP - CV_I) + CL_{IL}CIL + A_{transported_{OATP2B1}} \tag{S5}$$

3. OATP transporters on the basolateral membrane of hepatocytes mediate the uptake of the fu of PFOA from the extracellular space to the intracellular space Equation S6

$$\frac{dA_{L_{ec}}}{dt} = Q_I CV_I + Q_L C_P - (Q_I + Q_L)CV_{L_{ec}} - A_{transported_{OATP1B1,OATP1B3}} \tag{S6}$$

4. Finally, BSEP mediates the excretion of the fu of PFOA to the bile, at the same rate as its endogenous substrates, bile acids (de Bruijn et al., 2024), Equation S7.

$$\frac{dA_{L_{ic}}}{dt} = A_{transported_{OATP1B1,OATP1B3}} - A_{transported_{BSEP}} \tag{S7}$$

Even though physiologically the bile is stored in the gallbladder and is secreted to the intestinal lumen post-prandially, we assumed that it is continuously secreted to the intestinal lumen. We assumed this was correct for our modelling purposes which span over multiple years, minimizing the influence of hourly variations in elimination rates.

#### 1.6 Renal elimination and reabsorption

The two processes driving the renal elimination and reabsorption (RER) of PFOA are glomerular filtration and active reabsorption. To describe these processes, we developed a semi-permeability-limited, sequential kidney model, which is a simplified model from the fully permeability-limited sequential kidney models previously developed for drugs (Huang and Isoherranen, 2018; Pletz et al., 2021).

The final model equations for the kidney Equation S8, Equation S9, Equation S10, Equation S11, describe the following physiological and kinetic mechanisms:

1. The fu of PFOA in plasma is filtrated via the GFR to the primary urine in the proximal tubule lumen (PTL)

2. In the PTL, OAT4 transporter proteins mediate the reabsorption of the fu of PFOA in the proximal tubule lumen to the proximal tubular tissue (PTT).
3. The fraction of PFOA that is not filtrated follows the renal plasma flow into the PTT, from which it follows the same flow to reach the rest of kidney tissue (RKT) and thereafter the venous plasma.
4. The remaining PFOA in the PTL follows the proximal tubular fluid flow to the lumen of the rest of kidney (RKL) and subsequently to the urine.
5. PFOA is excreted in the urine based on physiological urine excretion rate.

$$\frac{dA_{PT_L}}{dt} = GFRfu_P C_P - Q_T C_{PT_L} - A_{transported_{OAT4}} \tag{S8}$$

$$\frac{dA_{PT_T}}{dt} = Q_K(C_P - C_{PT_T}) + A_{transported_{OAT4}} \tag{S9}$$

$$\frac{dA_{RK_T}}{dt} = Q_K(C_{PT_T} - CV_{RK_T}) \tag{S10}$$

$$\frac{dA_{RK_L}}{dt} = Q_T C_{PT_L} - Q_{Ur} C_{RK_L} \tag{S11}$$

We separated the proximal tubules in an explicit compartment were active reabsorption takes place. In initial model versions, we also explicitly modelled the distal tubules, and we lumped the loop of Henle and collecting ducts in a "mix" compartment. This complexity did not significantly alter the PFOA plasma concentration nor half-life predictions. As model complexity increases the model uncertainty and computational needs, we decided to simplify the model and lump all kidney sections other than the proximal tubules into a 'rest-of-kidney" compartment.

#### 1.7 Calculating the fraction unbound throughout lifetime

The fraction unbound of PFOA has been extensively studied; fu in plasma values derived using different techniques, ranging from the rapid equilibrium dialysis (RED) assay, ultra-centrifugation (UC) and solid phase microextraction (SPME), range from 0.00061 to 0.00245 (Fischer et al., 2024; Ryu et al., 2024; Smeltz et al., 2023). Given the high binding affinity of PFOA to plasma proteins, but also laboratory materials, most of the conventional methods available for deriving fu in plasma, such as the RED device, or UC, could be under-predicting the fu in plasma. Thereby, we considered that the sorption affinity of PFOA to albumin derived using the solid-phase micro extraction method would be the most accurate, given the method's capacity to concentrate the test compound before measuring it (Fischer et al., 2024). (Fischer et al., 2024) showed that both albumin and $\gamma$-globulin could be contributing to the binding of PFAS in plasma. However, for PFOA, albumin is the main determinant of fup, given the higher affinity of PFOA to albumin, compared to $\gamma$-globulin (Ryu et al., 2024; Fischer et al., 2025).

##### 1.8 Final model parameters

Table S2 and Table S3 contain an example of the final physiological and toxicokinetic parameters of the model, for a 30-year old woman. The columns contain the parameter, abbreviations as used in the model, unit, value and parameter reference.

#### 2 Results

##### 2.1 Model predictions

This study introduces a NG-PBK model for PFOA in humans, which accounts for physiological changes throughout lifetime, from birth to the elderly stages. In Figure S9 and Figure S10 are illustrated the results an adult, and of a simulated population representative of all life-stages (infants, toddlers, other children, teenagers, adults, elderly and very elderly) exposed to their respective reported daily median lower and upper bound dietary exposure(Panel), 2020). In Figure S9, the subject had a daily exposure of 0.19 ng/kg bw for 30 years. In Figure S10, the population had a daily exposure of one year of 0.19 or 7.23 ng/kg bw, followed by a wash-out period of 4 years.

It can be observed that all age groups had the same Cmax, and Tmax ( Figure S10). However, during the four years of no exposure (virtual wash-out period), the area under the concentration over time curve of PFOA was different between age groups, showing that the internal exposure was different. Following a four-year wash-out period, a significant decrease in serum concentration of PFOA was observed in infants and toddlers in comparison to the elderly. This observation can be attributed to variations in body weight during development, rather than to differences in the eliminating capacity. As the body grows during development, the total amount of PFOA in the body is virtually diluted. Figure S10 also shows that the predicted concentrations from either the highest or lowest bound exposures are close to the reported median plasma concentrations in children (3. 3 (0.49-6.9)) and adults (1. 9 (0.76-4.9)) ng/ml, indicated by the grey ribbon. This is an initial evaluation of whether the predictions are correct. A thorough evaluation of the observed versus predicted plasma concentrations, and half-lives is presented in the main article.

Both Figure S10 and Figure S11 illustrate the toxicokinetic differences between sexes, especially in the age range of 18- to 45-years old. Interestingly, this age range corresponds to the years when women actively menstruate. In these age groups, females were predicted to have lower plasma concentration compared to

Table S2: Model Physiological Parameters

| Parameter | Abbreviation | Unit | Initial_value | Reference |
|---|---|---|---|---|
| Body weight | BW | kg | 7.06e+01 | ICRP 89 |
| Cardiac output | QC | L/day | 4.99e+03 | ICRP 89 |
| Hematocrit | Hct | NA | 3.65e-01 | ICRP 89 |
| Fractional volume of intestine | VIc | NA | 1.60e-02 | ICRP 89 |
| Fractional volume of intestinal lumen | VILc | NA | 3.85e-02 | ICRP 89 |
| Surface area of the small intestine | SA_SIc | cm2 | 1.10e+03 | Calculated based on Willmann et al. 2004 |
| Fractional volume of liver | VLc | NA | 2.37e-02 | ICRP 89 |
| Fractional volume of live intracellular space | VL_icc | NA | 5.73e-01 | Calculated based on Utsey et al. 2020 |
| Fractional volume of liver extracellular space | VL_ecc | NA | 4.27e-01 | Calculated based on Utsey et al. 2020 |
| Fractional volume of kidney | VKc | NA | 4.60e-03 | ICRP 89 |
| Fractional volume of proximal tubule | VPTc | NA | 3.98e-01 | Calculated based on Pletz et al. 2020 |
| Fractional volume of proximal tubule tissue | VPTTc | NA | 5.47e-01 | Calculated based on Pletz et al. 2020 |
| Fractional volume of proximal tubule lumen | VPTLc | NA | 4.03e-01 | Calculated based on Pletz et al. 2020 |
| Fractional volume of rest of kidney lumen | VRKLc | NA | 2.83e-01 | Calculated based on Pletz et al. 2020 |
| Fractional volume of adipose | VAc | NA | 3.45e-01 | ICRP 89 |
| Adipose mass | MAc | NA | 7.68e+00 | ICRP 89 |
| Fractional volume of plasma | VPc | NA | 6.50e-02 | ICRP 89 |
| Fractional blood flow to the gut | QIc | NA | 1.65e-01 | ICRP 89 |
| Fractional blood flow to the liver | QLc | NA | 6.61e-02 | ICRP 89 |
| Fractional blood flow to the kidney | QKc | NA | 1.75e-01 | ICRP 89 |
| Fractional blood flow to the adipose | QAc | NA | 9.47e-02 | ICRP 89 |
| Fractional urine flow rate | QUrc | NA | 2.20e-02 | ICRP 89 |
| Glomerular filtration rate | GFR | NA | 1.66e+02 | ICRP 89 |
| Relative tubular fluid flow rate | QTc | L/day | 3.60e-01 | Scotcher et al. 2016; Pletz et al. 2020 |
| Colonic transit time | tco | /day | 3.36e+00 | Willmann et al. 2004 |
| Plasma to proximal tubule lumen albumin concentration ratio | R_PTL | NA | 2.99e+03 | Calculated based on Tojo and Satoshi 2012 |
| Plasma to liver albumin concentration ratio | R_L_ec | NA | 3.58e+07 | Calculated based on Utsey et al. 2020 |

Table S3: Model Toxicokinetic Parameters

| Parameter | Abbreviation | Unit | Initial_value | Reference |
|---|---|---|---|---|
| PFOA Fraction unbound in plasma | fup | NA | 5.22e-04 | Calculated, Fischer et al. 2024 |
| PFOA Partition coefficient to the intestine | PIc | NA | 6.90e+01 | Calculated based on Allendorf et al. 2021 |
| PFOA Partition coefficient to the liver | PLc | NA | 1.44e+02 | Calculated based on Allendorf et al. 2021 |
| PFOA Partition coefficient to the kidney | PKc | NA | 1.71e+02 | Calculated based on Allendorf et al. 2021 |
| PFOA Partition coefficient to the adipose | PAc | NA | 1.88e+02 | Calculated based on Allendorf et al. 2021 |
| PFOA Partition coefficient to the rest compartment | PRc | NA | 8.50e+01 | Calculated based on Allendorf et al. 2021 |
| PFOA Apparent permeability from the small intestine | Papp_SI | cm/s | 7.30e-06 | Janssen et al. 2024 |
| PFOA OATP2B1 Transporter maximum velocity | Vmax_OATP2B1c | umol/min/mg protein | 1.49e-03 | Lin et al.2023 |
| PFOA OATP2B1 Transporter affinity | Km_OATP2B1c | ug/L | 1.49e+02 | Lin et al.2023 |
| OATP2B1 IVIVE Scaling factor | SF_OATP2B1 | NA | 2.56e+05 | Calculated based on Lin et al.2023 |
| PFOA OATP1B1 Transporter maximum velocity | Vmax_OATP1B1c | umol/min/mg protein | 2.31e-03 | Lin et al.2023 |
| PFOA OATP1B1 Transporter affinity | Km_OATP1B1c | ug/L | 5.27e+01 | Lin et al.2023 |
| PFOA OATP1B3 Transporter maximum velocity | Vmax_OATP1B3c | umol/min/mg protein | 2.69e-03 | Lin et al.2023 |
| PFOA OATP1B3 Transporter affinity | Km_OATP1B3c | ug/L | 9.16e+01 | Lin et al.2023 |
| OATP1B1 IVIVE Scaling factor | SF_OATP1B1 | NA | 3.00e+06 | Calculated based on Lin et al.2023 |
| OATP1B3 IVIVE Scaling factor | SF_OATP1B3 | NA | 2.50e+05 | Calculated based on Lin et al.2023 |
| PFOA BSEP Transporter maximum velocity | Vmax_BSEPc | umol/min/mg BSEP | 7.00e+00 | Lin et al.2023 |
| PFOA BSEP Transporter affinity | Km_BSEPc | ug/L | 1.64e+01 | De Bruijn et al. 2024 |
| BSEP IVIVE Scaling factor | SF_BSEP | NA | 1.16e+01 | De Bruijn et al. 2024 |
| PFOA OAT4 Tranposter maximum velocity | Vmax_OAT4c | umol/min/mg protein | 4.50e-03 | Louisse et al. 2024 |
| PFOA OAT4 Transporter affinity | Km_OAT4c | ug/L | 4.70e+01 | Louisse et al. 2024 |
| OAT4 IVIVE Scaling factor | SF_OAT | NA | 5.13e+06 | Calculated based on Al-Madjoub et al 2021, and Lin et al. 2023 |
| Menstrual plasma clearance | CL_menses | L/day | 8.56e-04 | NA |
| Time to stop the exposure | expSTOP | day | 3.65e+02 | EFSA 2020 |
| Oral concentration | expOral | ug/kg/day | 4.18e-02 | NA |

males. Interestingly although in Figure S10 it can be observed that females seem to eliminate PFOA faster than men, these differences in elimination half-life were less observed in Figure S10. In Figure S10, it can be observed that both sexes have a relatively similar half-life.

Nevertheless, both plots ( Figure S10 and Figure S11) show the significance of accounting for physiological changes, even during adulthood. It can be observed that, in a simulated exposure and wash-out period throughout adulthood, the predicted half-life was longer when the physiological changes were accounted for. Furthermore, when physiological changes were accounted for, the plasma concentration continued to increase throughout the exposure duration, while in the simulation with default physiological values, it had reached steady-state within 10 years.

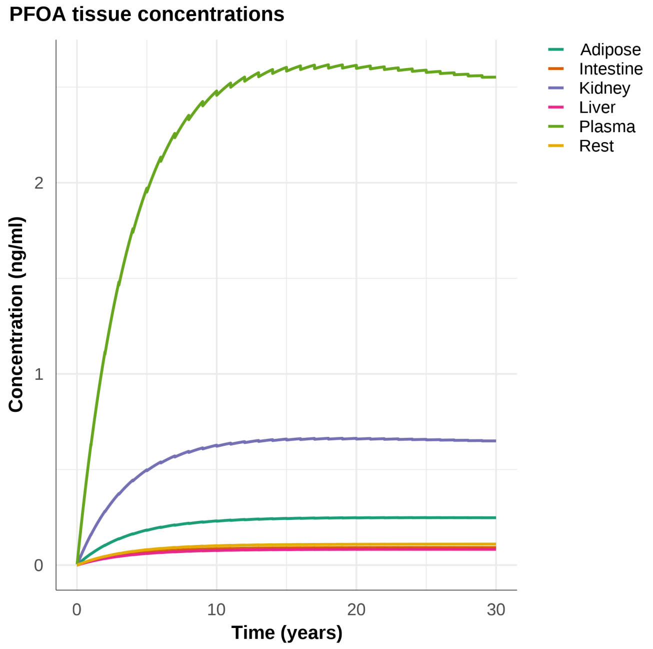


Figure S9: Predicted PFOA tissue concentrations (ng/ml) over time (years) in a man. The subject was simulated to have a daily exposure of PFAS of 0.19 ng/kg bw, corresponding to the median lower bound dietary exposure of adults. Exposure started at 20- and lasted until 50-years of age.

#### 2.2 Goodness-of-fit evaluation

The model was evaluated using the Olsen population, of 23 subjects (including 3 women) that were exposed to high PFOA doses due to occupational exposure (Olsen et al., 2007). In the original study, all subjects were recruited after

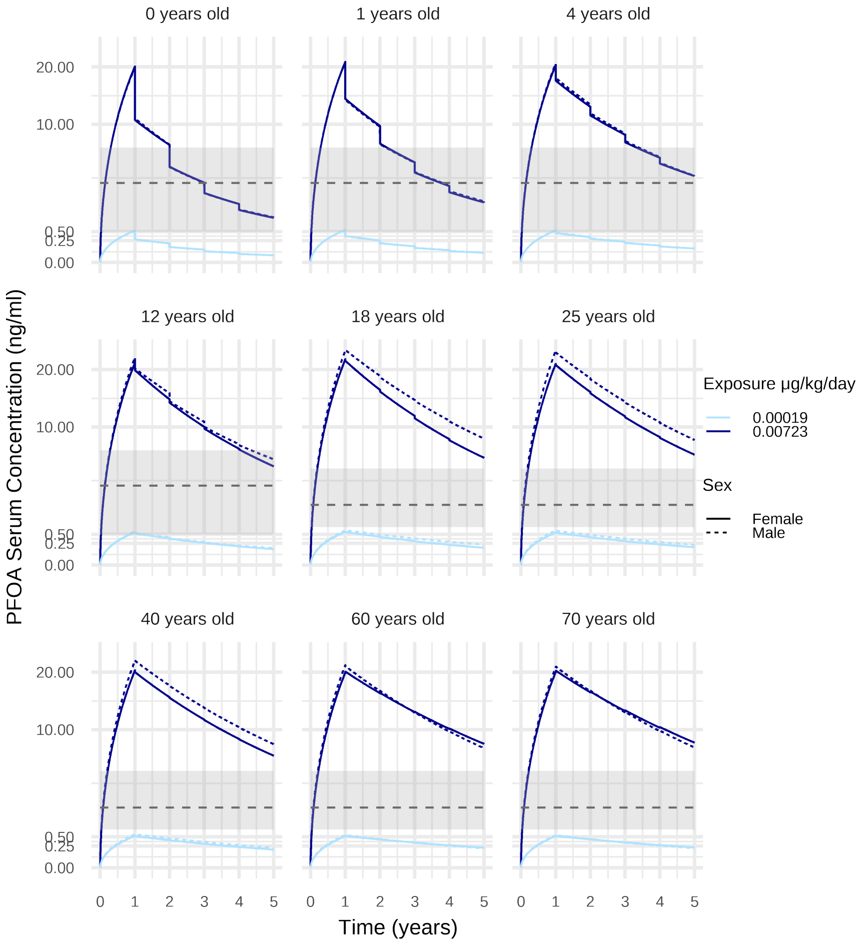


Figure S10: PFOA serum concentration (ng/ml) over time (years) in a population representative of all life stages, infants, toddlers, other children, teenagers, adults, elderly and very elderly. The population was simulated to have a daily dietary exposure of either 0.19 or 7.23 ng/kg bw for all age groups, which are the median of the reported median lower and upper bound dietary exposure of all age groups. The exposure lasted 1 year and was followed by a wash-out period of 4 years. Solid lines represent females and dashed lines males, the colour gradient of the lines (light to dark blue) represents the exposure (from low to high). The grey dotted line and ribbon illustrate the median plasma concentration per age group and its range, as reported in EFSA (3.3 (0.49-6.9) ng/ml in children and 1.9 (0.76-4.9) ng/ml in adults).

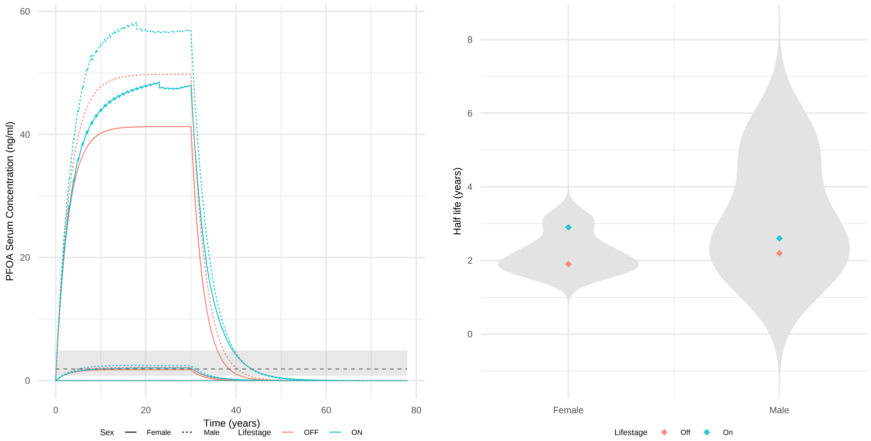


Figure S11: Model predictions of two adults, one male and one female, exposed to 0.18 ng/kg bw PFOA per day for 30 years, followed by a wash-out period of another 30 years. Their age at the start of exposure was 18 years old. In blue is a simulation with the lifestage option “On”, while in pink a simulation with the lifestage option “Off”, where “On” reflects a simulation where physiological changes over lifetime were taken into consideration and “Off” one where it was assumed that the subject had the reference BW of 70 kg and serum albumin concentration of 50 mg/mL throughout the whole simulation time. On the left: Predicted PFOA plasma concentration over time curve; solid and dashed lines represent female and male respectively; the grey dotted line and ribbon illustrate the median plasma concentration of 1.9 (0.76-4.9) ng/ml in adults). On the right: Predicted half-lives (in years) of PFOA plotted, against the distribution of half-lives reported in literature, grouped by sex; diamonds represent the predicted half-life per subject.

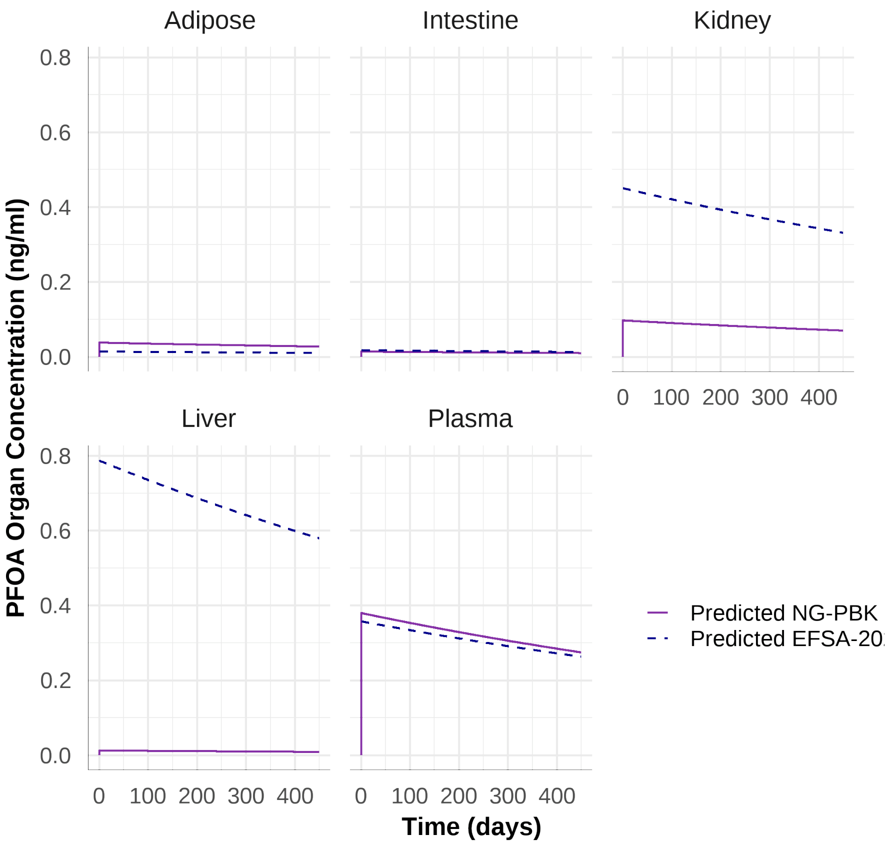


Figure S12: Evaluation of the NG-PBK model against the reference EFSA-2020 model (Panel), 2020). The plots compare the predicted PFOA concentration over time in the adipose, intestinal, kidney, liver and plasma tissues. Solid purple and dashed blue lines represent PFOA concentrations predicted by the NG-PBK and EFSA-2020 models respectively. Both models were run with the settings corresponding to the single oral $0.048\mu g/bw/day$exposure of the subject (adult male, 64 years old) in the Abraham study.

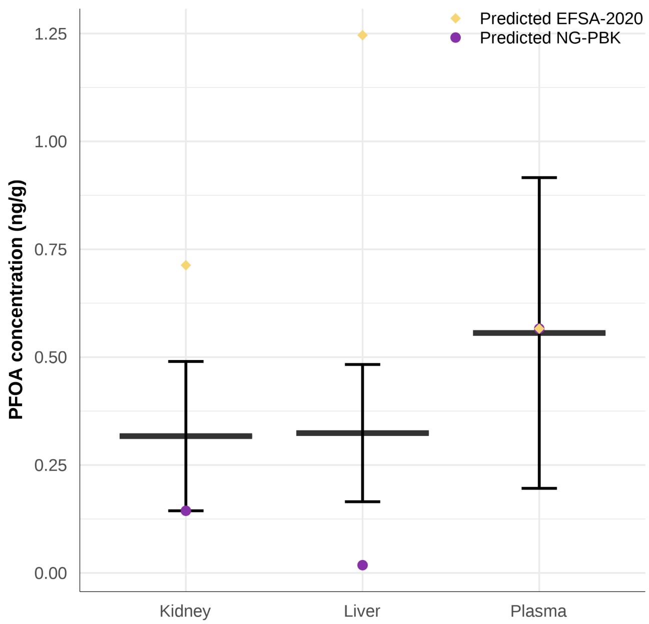


Figure S13: Comparison of the predicted and measured PFOA liver and kidney concentrations at a plasma concentration equal to that measured in human autopsies by (Nielsen et al., 2023). The daily oral exposure of twenty years, necessary to reach the measured blood concentrations by (Nielsen et al., 2023), was estimated by reverse dosimetry. A plasma:whole blood ratio of 2:1 was used to calculate the corresponding plasma concentrations. The solid lines and error-bars represent the measured concentrations, while in purple points are represented the NG-PBK model predicted concentrations and in yellow diamonds the EFSA-2020 model predicted concentrations.

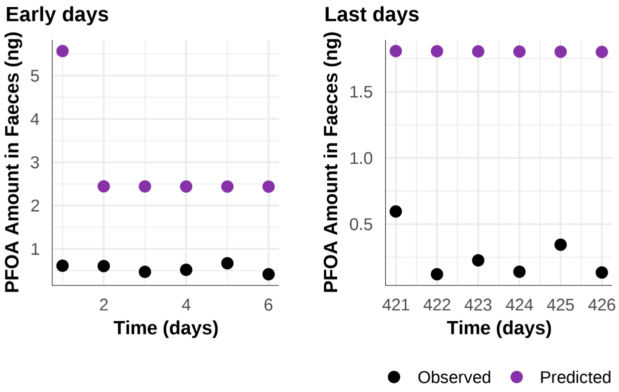


Figure S14: PFOA amount in feces excreted per day, at 1 to 6, and 421 to 426 days after oral intake. In dark points is the measured PFOA amount in feces from (Abraham et al., 2024) and in purple points the predicted one. Faecal clearance (mL/day/kg) was estimated by dividing the amount (ng) of PFOA in feces at a given day with the PFOA plasma concentration (ng/ml) on the respective day.

Table S4: Predicted and observed PFOA amount in feces, and fecal clearance. "P.A" and "O.A" stand for predicted and observed amount respectively (ng), "P.CL" and "O.CL" stand for predicted and observed daily fecal clearance (ml/day), "P.CL.bw" and "O.CL.bw" stand for predicted faecal clearance (ml/day/kg BW).

| Time (day) | P.A | O.A | P.CL | O.CL | P.CL.bw | O.CL.bw |
|---|---|---|---|---|---|---|
| 1 | 5.57 | 0.612 | 14.70 | 1.260 | 0.179 | 0.015 |
| 2 | 2.44 | 0.604 | 6.44 | 1.540 | 0.078 | 0.019 |
| 3 | 2.44 | 0.470 | 6.44 | 1.020 | 0.078 | 0.013 |
| 4 | 2.44 | 0.518 | 6.44 | 1.270 | 0.078 | 0.016 |
| 5 | 2.44 | 0.668 | 6.44 | 1.690 | 0.078 | 0.021 |
| 6 | 2.44 | 0.415 | 6.44 | 1.040 | 0.078 | 0.013 |
| 421 | 1.81 | 0.596 | 6.44 | 1.730 | 0.078 | 0.021 |
| 422 | 1.81 | 0.122 | 6.44 | 0.355 | 0.078 | 0.004 |
| 423 | 1.80 | 0.227 | 6.44 | 0.660 | 0.078 | 0.008 |
| 424 | 1.80 | 0.141 | 6.44 | 0.410 | 0.078 | 0.005 |
| 425 | 1.80 | 0.345 | 6.44 | 1.000 | 0.078 | 0.012 |
| 426 | 1.80 | 0.135 | 6.44 | 0.393 | 0.078 | 0.005 |
| Mean | 2.38 | 0.404 | 7.12 | 1.030 | 0.078 | 0.013 |
| Min | 1.80 | 0.122 | 6.44 | 0.355 | 0.078 | 0.004 |
| Max | 5.57 | 0.668 | 14.70 | 1.730 | 0.179 | 0.021 |

retirement from their work at the 3M factory, and PFOA plasma concentrations were measured twice, in order to determine the elimination half-life. The exact exposure route was not described in (Olsen et al., 2007), therefore in this step we mainly aimed to evaluate the capacity of the model to accurately predict the decline in PFOA serum concentration during the wash-out-phase. Physiological information of the population was estimated based on sex and age, while PFOA exposure was estimated via reverse dosimetry. Starting points for reverse dosimetry were PFOA serum concentration at the first sampling time (0.6 to 11.5 years after retirement), the exposure duration (assumed to be equal to the employment duration) and the exposure scenario (assumed to be dermal). Median estimated exposure was at 0.031 $\mu g/day/kgBW$, with a range of 0.005 to 0.397 $\mu g/day/kgBW$. Subjects were simulated to have dermal PFOA exposure throughout their employment and a wash-out period, without any PFOA exposure, until the time of their second serum sampling, which was also the end of the simulation time.

The model was also evaluated against three populations included in the study of (Johanson et al., 2023). The first group of the study included employees at the Arvidsjaur airport who were assumed to be exposed to PFOA throughout their whole employment time (Xu et al., 2020). The second and third groups included residents of Lulnaset and Visby, respectively, which were exposed to PFOA via their private water wells (Johanson et al., 2023). Given that more information regarding the exposure route and conditions for these populations was available, the aim of this simulation was to evaluate whether the NG-PBK model could accurately predict the measured plasma concentrations, based on the given exposure estimations.

Similar to the Olsen population, physiological information for the populations was estimated based on sex and age. PFOA exposure was estimated based on the measured PFOA concentrations in drinking water at the three locations, and an assumed consumption of 2L of water per day, except for Arvidsjaur employees being only exposed to PFOA via drinking water during work time adjusted by half of the daily consumption. A background exposure via food consumption was also included and assumed to be at 0.00018 $\mu g/kg/day$, corresponding to the median lower bound dietary exposure (Panel), 2020).

Subjects were simulated to have an oral PFOA exposure throughout their employment (Arvidsjaur), or residence (Lulnaset and Visby) time and a wash-out period, without any PFOA exposure, until the time of serum sampling, which was also the end of the simulation time. Finally, Predicted PFOA serum concentrations were compared to those measured (Johanson et al., 2023).

#### 2.3 Global Sensitivity Analysis

##### 2.3.1 Morris Sensitivity Analysis

Contrary to our results, an epidemiological study from a highly exposed population in Italy showed that age had a stronger correlation to PFOA half-life

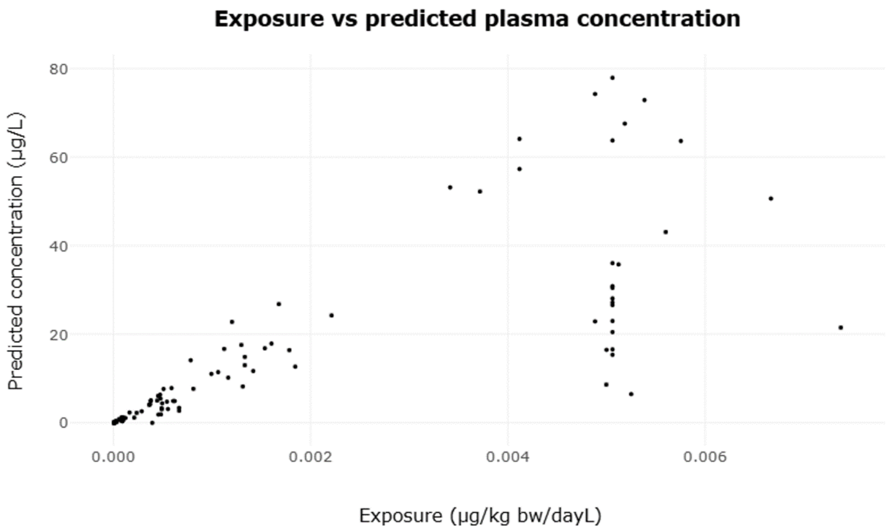


Figure S15: Estimated exposure Vs Observed Serum concentration in all the Swedish populations together.

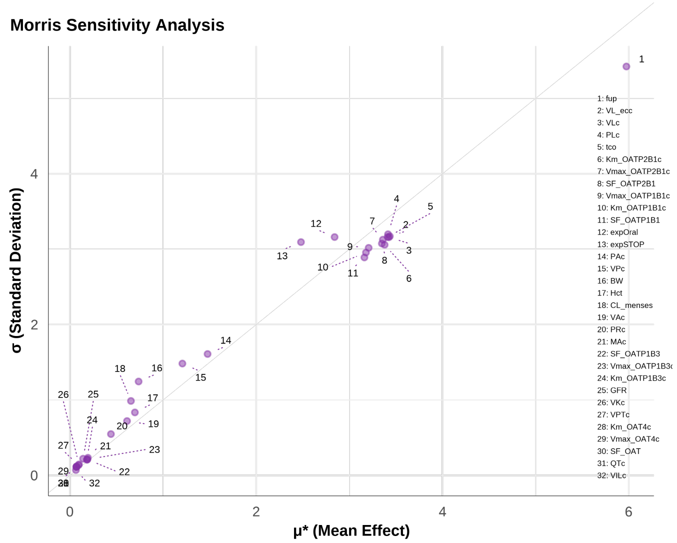


Figure S16: Results of the Morris test on all model outputs together

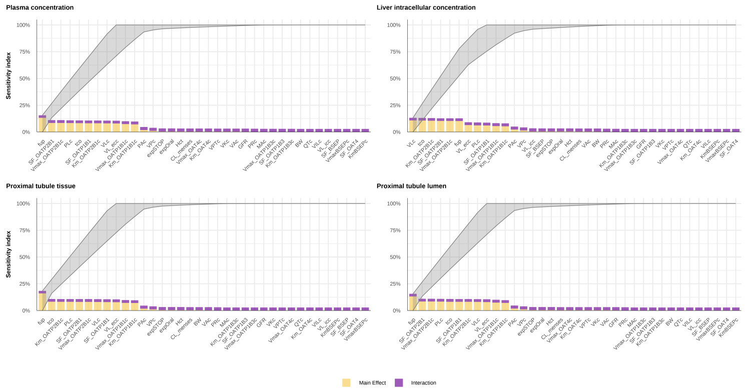


Figure S17: Results of the eFAST test on model outputs; A: PFOA plasma concentration, B: PFOA proximal tubule tissue concentration, C: PFOA liver intracellular space concentration

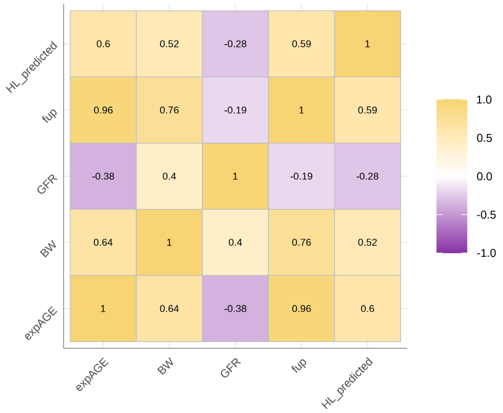


Figure S18: Results of the Pearson correlation analysis of the male subjects aged from 0- to 64-years). The plot shows the correlation between physiological parameters (age at exposure (expAGE), body weight (BW), GFR, fu in plasma (fup) (dependent on the differences in serum albumin concentration over lifetime), and PFOA clearance through menstruation (CL_menses)) with the predicted half-life of PFOA (HL_predicted). The higher the correlation the darker the colour becomes, with the yellow colour illustrating a positive correlation and the purple colour a negative correlation.

compared to GFR (Batzella et al., 2024). However, we believe that the study was not able to show the high correlation between GFR and half-life because the age range of the participants was small and within the age range (18.5 to 36.7) where no major changes in GFR happen. As shown in Figure S2, GFR increases until 18 years of age, then reaches an almost steady state until 40 years of age, from when an exponential decline in GFR happens (Smeets et al., 2022; Astley et al., 2025).

These results are in agreement with previous studies, where fup was found to be the most influencing parameter to the model's output (Fischer et al., 2025; Loccisano et al., 2011; Husøy et al., 2023; Lin et al., 2023; Panel), 2020; Ratier et al., 2024). Contrary to previously published PBK models, where fup was fitted to animal data (Loccisano et al., 2011; Husøy et al., 2023; Lin et al., 2023; Panel), 2020; Ratier et al., 2024), the fup in our model was calculated based on the binding affinity of PFOA to albumin and $\gamma$-globulin, and the concentration of albumin and $\gamma$-globulin in plasma.

In our model we did not include pregnancy, nor breast feeding, both of which are known elimination routes of PFOA in women (Panel), 2020; Batzella et al., 2024). These excretion routes should be included when modelling specifically PFOA toxicokinetics during pregnancy and breast feeding, or when one is interested in PFOA toxicokinetics in foetus and infants.

### References

Abraham, K., Mertens, H., Richter, L., Mielke, H., Schwerdtle, T., Monien, B.H., 2024. Kinetics of 15 per- and polyfluoroalkyl substances (pfas) after single oral application as a mixture – a pilot investigation in a male volunteer. Environment International 193, 109047. URL: http://dx.doi.org/10.1016/j.envint.2024.109047, doi:10.1016/j.envint.2024.109047.

Allendorf, F., Berger, U., Goss, K.U., Ulrich, N., 2019. Partition coefficients of four perfluoroalkyl acid alternatives between bovine serum albumin (bsa) and water in comparison to ten classical perfluoroalkyl acids. Environmental Science: Processes & Impacts 21, 1852–1863. URL: http://dx.doi.org/10.1039/C9EM00290A, doi:10.1039/C9EM00290A.

Allendorf, F., Goss, K.U., Ulrich, N., 2021. Estimating the equilibrium distribution of perfluoroalkyl acids and 4 of their alternatives in mammals. Environmental Toxicology and Chemistry 40, 910–920. URL: https://setac.onlinelibrary.wiley.com/doi/abs/10.1002/etc.4954, doi:10.1002/etc.4954.

Astley, M.E., Chesnaye, N.C., Hallan, S., Gambaro, G., Ortiz, A., Carrero, J.J., Ebert, N., Eriksen, B., Faucon, A.L., Ferraro, P.M., Indridason, O.S., Ittermann, T., Jonsson, A.J., Rise Langlo, K., Melsom, T., Schaeffner, E., Stracke, S., Stel, V.S., Jager, K.J., 2025. Age- and sex-specific reference values of estimated glomerular filtration rate for european adults. Kidney

International 107, 1076–1087. URL: http://dx.doi.org/10.1016/j.kint.2025.02.025, doi:10.1016/j.kint.2025.02.025. citation Key: astley2025.

Batzella, E., Rosato, I., Pitter, G., Da Re, F., Russo, F., Canova, C., Fletcher, T., 2024. Determinants of pfoa serum half-life after end of exposure: A longitudinal study on highly exposed subjects in the veneto region. Environmental Health Perspectives 132. URL: http://dx.doi.org/10.1289/EHP13152, doi:10.1289/ehp13152.

de Bruijn, V., te Kronnie, W., Rietjens, I.M.C.M., Bouwmeester, H., 2024. Intestinal in vitro transport assay combined with physiologically based kinetic modeling as a tool to predict bile acid levels in vivo. ALTEX - Alternatives to animal experimentation 41, 20–36. URL: http://dx.doi.org/10.14573/altex.2302011, doi:10.14573/altex.2302011. citation Key: de_Bruijn_te2024.

Cao, H., Zhou, Z., Hu, Z., Wei, C., Li, J., Wang, L., Liu, G., Zhang, J., Wang, Y., Wang, T., Liang, Y., 2022. Effect of enterohepatic circulation on the accumulation of per- and polyfluoroalkyl substances: Evidence from experimental and computational studies. Environmental Science & Technology 56, 3214–3224. URL: http://dx.doi.org/10.1021/acs.est.1c07176, doi:10.1021/acs.est.1c07176. citation Key: cao2022.

Ebert, A., Allendorf, F., Berger, U., Goss, K.U., Ulrich, N., 2020. Membrane/water partitioning and permeabilities of perfluoroalkyl acids and four of their alternatives and the effects on toxicokinetic behavior. Environmental Science & Technology 54, 5051–5061. URL: https://doi.org/10.1021/acs.est.0c00175, doi:10.1021/acs.est.0c00175.

Fischer, F.C., Ludtke, S., Thackray, C., Pickard, H.M., Haque, F., Dassuncao, C., Endo, S., Schaider, L., Sunderland, E.M., 2024. Binding of per- and polyfluoroalkyl substances (pfas) to serum proteins: Implications for toxicokinetics in humans. Environmental Science & Technology 58, 1055–1063. URL: https://doi.org/10.1021/acs.est.3c07415, doi:10.1021/acs.est.3c07415.

Fischer, F.C., Thackray, C., Ferguson, N., Chicoine, C., Skende, O., Hu, Z., Zhu, Y., Slitt, A., Sunderland, E.M., 2025. Understanding mechanisms of pfas absorption, distribution, and elimination using a physiologically based toxicokinetic model. Environmental Science & Technology 59, 13240–13250. URL: https://pubs.acs.org/doi/10.1021/acs.est.5c05473, doi:10.1021/acs.est.5c05473.

Huang, W., Isoherranen, N., 2018. Development of a dynamic physiologically based mechanistic kidney model to predict renal clearance. CPT: Pharmacometrics & Systems Pharmacology 7, 593–602. URL: https://ascpt.onlinelibrary.wiley.com/doi/10.1002/psp4.12321, doi:10.1002/psp4.12321.

Husøy, T., Caspersen, I., Thépaut, E., Knutsen, H., Haug, L., Andreassen, M., Gkrillas, A., Lindeman, B., Thomsen, C., Herzke, D., Dirven, H., Wojewodzic, M., 2023. Comparison of aggregated exposure to perfluorooctanoic acid (pfoa) from diet and personal care products with concentrations in blood using a pbpk model – results from the norwegian biomonitoring study in euromix. Environmental Research 239, 117341. URL: http://dx.doi.org/10.1016/j.envres.2023.117341, doi:10.1016/j.envres.2023.117341. citation Key: husøy2023.

Janssen, A.W.F., Duivenvoorde, L.P.M., Beekmann, K., Pinckaers, N., Van Der Hee, B., Noorlander, A., Leenders, L.L., Louisse, J., Van Der Zande, M., 2024. Transport of perfluoroalkyl substances across human induced pluripotent stem cell-derived intestinal epithelial cells in comparison with primary human intestinal epithelial cells and caco-2 cells. Archives of Toxicology 98, 3777–3795. URL: https://link.springer.com/10.1007/s00204-024-03851-x, doi:10.1007/s00204-024-03851-x.

Johanson, G., Gyllenhammar, I., Ekstrand, C., Pyko, A., Xu, Y., Li, Y., Norström, K., Lilja, K., Lindh, C., Benskin, J.P., Georgelis, A., Forsell, K., Jakobsson, K., Glynn, A., Vogs, C., 2023. Quantitative relationships of perfluoroalkyl acids in drinking water associated with serum concentrations above background in adults living near contamination hotspots in sweden. Environmental Research 219, 115024. URL: http://dx.doi.org/10.1016/j.envres.2022.115024, doi:10.1016/j.envres.2022.115024.

Karakoltzidis, A., Karakitsios, S.P., Gabriel, C., Sarigiannis, D., 2025. Integrated pbpk modelling for pfoa exposure and risk assessment. Environmental Research 282, 121947. URL: http://dx.doi.org/10.1016/j.envres.2025.121947, doi:10.1016/j.envres.2025.121947. citation Key: karakoltzidis2025.

Kimura, O., Fujii, Y., Haraguchi, K., Kato, Y., Ohta, C., Koga, N., Endo, T., 2017. Uptake of perfluorooctanoic acid by caco-2 cells: Involvement of organic anion transporting polypeptides. Toxicology Letters 277, 18–23. URL: https://linkinghub.elsevier.com/retrieve/pii/S0378427417301893, doi:10.1016/j.toxlet.2017.05.012.

Kudo, N., Sakai, A., Mitsumoto, A., Hibino, Y., Tsuda, T., Kawashima, Y., 2007. Tissue distribution and hepatic subcellular distribution of perfluorooctanoic acid at low dose are different from those at high dose in rats. Biological and Pharmaceutical Bulletin 30, 1535–1540. URL: http://www.jstage.jst.go.jp/article/bpb/30/8/30_8_1535/_article, doi:10.1248/bpb.30.1535.

Lin, X., Xing, Y., Chen, H., Zhou, Y., Zhang, X., Liu, P., Li, J., Lee, H.K., Huang, Z., 2023. Characteristic and health risk of per- and polyfluoroalkyl substances from cosmetics via dermal exposure. Environmental Pollution 338, 122685. URL: https://linkinghub.elsevier.com/retrieve/pii/S0269749123016871, doi:10.1016/j.envpol.2023.122685.

Loccisano, A.E., Campbell, J.L., Andersen, M.E., Clewell, H.J., 2011. Evaluation and prediction of pharmacokinetics of pfoa and pfos in the monkey and human using a pbpk model. Regulatory Toxicology and Pharmacology 59, 157–175. URL: https://linkinghub.elsevier.com/retrieve/pii/S0273230010002242, doi:10.1016/j.yrtph.2010.12.004.

Louisse, J., Dellafiora, L., van den Heuvel, J.J.M.W., Rijkers, D., Leenders, L., Dorne, J.L.C.M., Punt, A., Russel, F.G.M., Koenderink, J.B., 2022. Perfluoroalkyl substances (pfass) are substrates of the renal human organic anion transporter 4 (oat4). Archives of Toxicology 97, 685–696. URL: http://dx.doi.org/10.1007/s00204-022-03428-6, doi:10.1007/s00204-022-03428-6. citation Key: louisse2022.

Louisse, J., Pedroni, L., van den Heuvel, J.J.M.W., Rijkers, D., Leenders, L., Noorlander, A., Punt, A., Russel, F.G.M., Koenderink, J.B., Dellafiora, L., 2024. In vitro and in silico characterization of the transport of selected perfluoroalkyl carboxylic acids and perfluoroalkyl sulfonic acids by human organic anion transporter 1 (oat1), oat2 and oat3. Toxicology 509, 153961. URL: https://www.sciencedirect.com/science/article/pii/S0300483X24002427, doi:10.1016/j.tox.2024.153961.

Maestri, L., Negri, S., Ferrari, M., Ghittori, S., Fabris, F., Danesino, P., Imbriani, M., 2006. Determination of perfluorooctanoic acid and perfluorooctanesulfonate in human tissues by liquid chromatography/single quadrupole mass spectrometry. Rapid Communications in Mass Spectrometry 20, 2728–2734. URL: https://analyticalsciencejournals.onlinelibrary.wiley.com/doi/10.1002/rcm.2661, doi:10.1002/rcm.2661.

Nielsen, F., Fischer, F.C., Leth, P.M., Grandjean, P., 2023. Occurrence of major perfluorinated alkylate substances in human blood and target organs. Environmental Science & Technology 58, 143–149. URL: http://dx.doi.org/10.1021/acs.est.3c06499, doi:10.1021/acs.est.3c06499. citation Key: nielsen2023.

Nigam, S.K., Granados, J.C., 2023. Oat, oatp, and mrp drug transporters and the remote sensing and signaling theory. Annual Review of Pharmacology and Toxicology 63, 637–660. URL: http://dx.doi.org/10.1146/annurev-pharmtox-030322-084058, doi:10.1146/annurev-pharmtox-030322-084058. citation Key: nigam2023.

Niu, S., Cao, Y., Chen, R., Bedi, M., Sanders, A.P., Ducatman, A., Ng, C., 2023. A state-of-the-science review of interactions of per- and polyfluoroalkyl substances (pfas) with renal transporters in health and disease: Implications for population variability in pfas toxicokinetics. Environmental Health Perspectives 131. URL: http://dx.doi.org/10.1289/EHP11885, doi:10.1289/ehp11885. citation Key: niu2023.

Olsen, G.W., Burris, J.M., Ehresman, D.J., Froehlich, J.W., Seacat, A.M., Butenhoff, J.L., Zobel, L.R., 2007. Half-life of serum elimination of perfluorooctanesulfonate,perfluorohexanesulfonate, and perfluorooctanoate in retired fluorochemical production workers. Environmental Health Perspectives 115, 1298–1305. URL: http://dx.doi.org/10.1289/ehp.10009, doi:10.1289/ehp.10009.

Panel), E.C., 2020. Risk to human health related to the presence of perfluoroalkyl substances in food. EFSA Journal 18. URL: https://data.europa.eu/

doi/10.2903/j.efsa.2020.6223, doi:10.2903/j.efsa.2020.6223. citation Key: EFSA2020.

Peyret, T., Poulin, P., Krishnan, K., 2010. A unified algorithm for predicting partition coefficients for pbpk modeling of drugs and environmental chemicals. Toxicology and Applied Pharmacology 249, 197–207. URL: https://linkinghub.elsevier.com/retrieve/pii/S0041008X10003443, doi:10.1016/j.taap.2010.09.010.

Pletz, J., Allen, T.J., Madden, J.C., Cronin, M.T., Webb, S.D., 2021. A mechanistic model to study the kinetics and toxicity of salicylic acid in the kidney of four virtual individuals. Computational Toxicology 19, 100172. URL: http://dx.doi.org/10.1016/j.comtox.2021.100172, doi:10.1016/j.comtox.2021.100172. citation Key: pletz2021.

Pottel, H., Hoste, L., Dubourg, L., Ebert, N., Schaeffner, E., Eriksen, B., Melsom, T., Lamb, E.J., Rule, A.D., Turner, S.T., Glassock, R.J., De Souza, V., Selistre, L., Mariat, C., Martens, F., Delanaye, P., 2016. An estimated glomerular filtration rate equation for the full age spectrum. Nephrology Dialysis Transplantation 31, 798–806. URL: https://academic.oup.com/ndt/article-lookup/doi/10.1093/ndt/gfv454, doi:10.1093/ndt/gfv454.

Poulin, P., Schoenlein, K., Theil, F., 2001. Prediction of adipose tissue: Plasma partition coefficients for structurally unrelated drugs. Journal of Pharmaceutical Sciences 90, 436–447. URL: https://linkinghub.elsevier.com/retrieve/pii/S0022354916307407, doi:10.1002/1520-6017(200104)90:4<436::AID-JPS1002>3.0.CO;2-P.

Ragnarsdóttir, O., Abou-Elwafa Abdallah, M., Harrad, S., 2024. Dermal bioavailability of perfluoroalkyl substances using in vitro 3d human skin equivalent models. Environment International 188, 108772. URL: http://dx.doi.org/10.1016/j.envint.2024.108772, doi:10.1016/j.envint.2024.108772. citation Key: ragnarsdóttir2024.

Ratier, A., Casas, M., Grazuleviciene, R., Slama, R., Småstuen Haug, L., Thomsen, C., Vafeiadi, M., Wright, J., Zeman, F.A., Vrijheid, M., Brochot, C., 2024. Estimating the dynamic early life exposure to pfoa and pfos of the helix children: Emerging profiles via prenatal exposure, breastfeeding, and diet. Environment International 186, 108621. URL: http://dx.doi.org/10.1016/j.envint.2024.108621, doi:10.1016/j.envint.2024.108621. citation Key: ratier2024.

Rodgers, T., Leahy, D., Rowland, M., 2005. Physiologically based pharmacokinetic modeling 1: Predicting the tissue distribution of moderate-to-strong bases. Journal of Pharmaceutical Sciences 94, 1259–1276. URL: https://linkinghub.elsevier.com/retrieve/pii/S0022354916317890, doi:10.1002/jps.20322.

Rodgers, T., Rowland, M., 2006. Physiologically based pharmacokinetic modelling 2: Predicting the tissue distribution of acids, very weak bases, neutrals and zwitterions. Journal of Pharmaceutical Sciences 95, 1238–1257. URL: https://linkinghub.elsevier.com/retrieve/pii/S0022354916320342, doi:10.1002/jps.20502.

Rodgers, T., Rowland, M., 2007. Mechanistic approaches to volume of distribution predictions: Understanding the processes. Pharmaceutical Research 24, 918–933. URL: https://link.springer.com/10.1007/s11095-006-9210-3, doi:10.1007/s11095-006-9210-3.

Ruggiero, M.J., Miller, H., Idowu, J.Y., Zitzow, J.D., Chang, S.C., Hagenbuch, B., 2021. Perfluoroalkyl carboxylic acids interact with the human bile acid transporter ntcp. Livers 1, 221–229. URL: https://www.mdpi.com/2673-4389/1/4/17, doi:10.3390/livers1040017.

Ryu, S., Yamaguchi, E., Sadegh Modaresi, S.M., Agudelo, J., Costales, C., West, M.A., Fischer, F., Slitt, A.L., 2024. Evaluation of 14 pfas for permeability and organic anion transporter interactions: Implications for renal clearance in humans. Chemosphere 361, 142390. URL: https://www.sciencedirect.com/science/article/pii/S0045653524012839, doi:10.1016/j.chemosphere.2024.142390.

Smeets, N.J., IntHout, J., van der Burgh, M.J., Schwartz, G.J., Schreuder, M.F., de Wildt, S.N., 2022. Maturation of gfr in term-born neonates: An individual participant data meta-analysis. Journal of the American Society of Nephrology 33, 1277–1292. URL: http://dx.doi.org/10.1681/ASN.2021101326, doi:10.1681/asn.2021101326. citation Key: smeets2022.

Smeltz, M., Wambaugh, J.F., Wetmore, B.A., 2023. Plasma protein binding evaluations of per- and polyfluoroalkyl substances for category-based toxicokinetic assessment. Chemical Research in Toxicology 36, 870–881. URL: https://doi.org/10.1021/acs.chemrestox.3c00003, doi:10.1021/acs.chemrestox.3c00003.

Tiburtini, G.A., Bertarini, L., Bersani, M., Dragani, T.A., Rolando, B., Binello, A., Barge, A., Spyrakis, F., 2024. In silico prediction of the interaction of legacy and novel per- and poly-fluoroalkyl substances (pfas) with selected human transporters and of their possible accumulation in the human body. Archives of Toxicology 98, 3035–3047. URL: https://link.springer.com/10.1007/s00204-024-03797-0, doi:10.1007/s00204-024-03797-0.

Utsey, K., Gastonguay, M.S., Russell, S., Freling, R., Riggs, M.M., Elmokadem, A., 2020. Quantification of the impact of partition coefficient prediction methods on physiologically based pharmacokinetic model output using a standardized tissue composition. Drug Metabolism and Disposition 48, 903–916. URL: http://dx.doi.org/10.1124/dmd.120.090498, doi:10.1124/dmd.120.090498. citation Key: Utsey2020b.

Valentin, J., 2002. Basic anatomical and physiological data for use in radiological protection: reference values. Annals of the ICRP 32, 1–277. URL: http://dx.doi.

org/10.1016/S0146-6453(03)00002-2, doi:10.1016/s0146-6453(03)00002-2. citation Key: ICRP89.

Vujic, E., Ferguson, S.S., Brouwer, K.L.R., 2024. Effects of pfas on human liver transporters: implications for health outcomes. Toxicological Sciences 200, 213–227. URL: https://academic.oup.com/toxsci/article/200/2/213/7667880, doi:10.1093/toxsci/kfae061.

Weaving, G., Batstone, G.F., Jones, R.G., 2015. Age and sex variation in serum albumin concentration: an observational study. Annals of Clinical Biochemistry: International Journal of Laboratory Medicine 53, 106–111. URL: http://dx.doi.org/10.1177/0004563215593561, doi:10.1177/0004563215593561. citation Key: weaving2015.

Xu, Y., Fletcher, T., Pineda, D., Lindh, C.H., Nilsson, C., Glynn, A., Vogs, C., Norström, K., Lilja, K., Jakobsson, K., Li, Y., 2020. Serum half-lives for short- and long-chain perfluoroalkyl acids after ceasing exposure from drinking water contaminated by firefighting foam. Environmental Health Perspectives 128. URL: http://dx.doi.org/10.1289/ehp6785, doi:10.1289/ehp6785.

Yang, C.H., Glover, K.P., Han, X., 2010. Characterization of cellular uptake of perfluorooctanoate via organic anion-transporting polypeptide 1a2, organic anion transporter 4, and urate transporter 1 for their potential roles in mediating human renal reabsorption of perfluorocarboxylates. Toxicological Sciences 117, 294–302. URL: https://academic.oup.com/toxsci/article-lookup/doi/10.1093/toxsci/kfq219, doi:10.1093/toxsci/kfq219.

Zhao, L., Teng, M., Zhao, X., Li, Y., Sun, J., Zhao, W., Ruan, Y., Leung, K.M., Wu, F., 2023. Insight into the binding model of per- and polyfluoroalkyl substances to proteins and membranes. Environment International 175, 107951. URL: http://dx.doi.org/10.1016/j.envint.2023.107951, doi:10.1016/j.envint.2023.107951. citation Key: zhao2023.
